# Deployment of a *Vibrio cholerae* ordered transposon mutant library in a quorum-competent genetic background

**DOI:** 10.1101/2023.10.31.564941

**Authors:** Nkrumah A. Grant, Gracious Yoofi Donkor, Jordan T. Sontz, William Soto, Christopher M. Waters

## Abstract

*Vibrio cholerae*, the causative agent of cholera, has sparked seven pandemics in recent centuries, with the current one being the most prolonged. *V. cholerae’s* pathogenesis hinges on its ability to switch between low and high cell density gene regulatory states, enabling transmission between host and the environment. Previously, a transposon mutant library for *V. cholerae* was created to support investigations aimed toward uncovering the genetic determinants of its pathogenesis. However, subsequent sequencing uncovered a mutation in the gene *luxO* of the parent strain, rendering mutants unable to exhibit high cell density behaviors. In this study, we used chitin-independent natural transformation to move transposon insertions from these low cell density mutants into a wildtype genomic background. Library transfer was aided by a novel gDNA extraction we developed using thymol, which also showed high lysis-specificity for *Vibrio*. The resulting Grant Library comprises 3,102 unique transposon mutants, covering 79.8% of *V. cholerae’s* open reading frames. Whole genome sequencing of randomly selected mutants demonstrates 100% precision in transposon transfer to cognate genomic positions of the recipient strain. Notably, in no instance did the *luxO* mutation transfer into the wildtype background. Our research uncovered density-dependent epistasis in growth on inosine, an immunomodulatory metabolite secreted by gut bacteria that is implicated in enhancing gut barrier functions. Additionally, Grant Library mutants retain the plasmid that enables rapid, scarless genomic editing. In summary, the Grant Library reintroduces organismal relevant genetic contexts absent in the low cell density locked library equivalent.

**Significance Statement:** Ordered transposon mutant libraries are essential tools for catalyzing research by providing access to null mutants of all non-essential genes. Such a library was previously generated for *Vibrio cholerae*, but whole genome sequencing revealed that this library was made using a parent strain that is unable to exhibit cell-cell communication known as quorum sensing. Here, we utilize natural competence combined with a novel, high-throughput genomic DNA extraction method to regenerate the signaling incompetent *V. cholerae* ordered transposon mutant library in quorum sensing competent strain. Our library provides researchers with a powerful tool to understand *V. cholerae* biology within a genetic context that influences how it transitions from an environmentally benign organism to a disease-causing pathogen.

## Introduction

*Vibrio cholerae* is a human pathogen and the causative agent of the acute diarrheal disease cholera. While cholera is a global threat, the disease is endemic in developing countries, where it is estimated that there are between 1.3 to 4.0 million *V. cholerae* annual infections, resulting in 21,000 – 143,000 deaths (1, 2). Implementation of disease intervention strategies, including increasing access to clean drinking water, provision of adequate healthcare infrastructure, and educating citizens in communities at risk of disease spread, remains a top priority in the fight toward eradicating cholera. As such, the Global Task Force on Cholera Control was founded in 2017 to support efforts aimed at reducing cholera burden by 90% before the year 2030 (3, 4).

To date there have been seven cholera pandemics. The seventh pandemic began in 1961 and is ongoing, marking it as the longest running cholera pandemic (5). *V. cholerae* strains are grouped by surface antigens, including O polysaccharide, such that there are >200 serogroups (6). Of these serogroups, only two, O1 and O139, host pandemic *V. cholerae* strains (7, 8). Furthermore, pandemic strains from the O1 serogroup are further stratified into two biotypes with the fifth and sixth pandemics caused by the Classical and the seventh caused by El Tor (9).

Like most vibrios, *V. cholerae* predominantly possesses two chromosomes (10, 11). Whole genome sequencing of strains before and after the emergence of seventh pandemic *V. cholerae* strains has proved to be an invaluable resource toward understanding its origin and evolution. These analyses show *V. cholerae* El Tor originated from a nonpathogenic strain (12), becoming a pathogen after acquiring two genomic islands (VSP-I and VSP-II), a prophage that expresses cholera toxin (CTXφ), and 12 additional mutations that enhanced its transmissibility (9, 13, 14). Recent work has also shown that these genomic changes contribute to seventh pandemic *V. cholerae’s* long-term persistence in the estuarian environments they occupy (15).

Key to El Tor’s transition to a pathogen is that it is naturally competent, granting it the ability to take up DNA from the environment which can be integrated into the genome by homologous recombination (16). Natural competence is a highly regulated process, and four environmental factors are key determinants for driving this process. These include nutrient limitation, extracellular signaling molecules, high cell density, and the presence of chitin (17–19), which together coordinate the expression of several regulators governing competence (20–23). Recent work has shown that expressing the master regulator of competence TfoX can circumvent the chitin requirement for natural competence in *V. cholerae* (19, 24). This discovery has allowed exploitation of chitin-independent natural competence as a genetic tool for editing *Vibrio spp.* genomes (25–27), facilitating our understanding of the functional relationships between genes and furthering our understanding of how *V. cholerae* interfaces with its environment.

Our understanding of *V cholerae’s* ecology and evolution has also benefited from transposon mutagenesis. In contrast to conventionally laborious genetic engineering approaches for disrupting single genes to study their phenotypic effects, transposon mutagenesis allows the simultaneous construction and screening of tens of thousands of mutants. Researchers have taken advantage of transposon mutagenesis to construct nonredundant (ordered) mutant libraries, where each nonessential gene has been inactivated and the individual mutants are arrayed across the wells of 96-well microtiter plates (28–31). An ordered mutant *V. cholerae* library was constructed in 2008 (32). The ordered library consists of >3,100 mutants, each with a Tn5-based transposon insertion that confers kanamycin resistance. The mutant library has provided a public resource enhancing *V. cholerae* genotypic and phenotypic investigations, supporting identifying novel therapeutic targets for vaccine development (33), and increasing our understanding of the *V. cholerae* virulome (34, 35), amongst other discoveries (36, 37).

Although significant research advances have been made in our understanding of *V. cholerae* using mutants from this ordered mutant transposon library, the library was constructed in a laboratory acquired quorum sensing mutant of *V. cholerae*. Quorum sensing, the process of cell-to-cell communication in bacteria, allows cells to undergo global changes in gene expression as the bacteria transition from low to high cell density. This transition can influence behaviors like host immunity escape (38), virulence factor production (39–41), biofilm formation (42), infectivity and environmental dissemination (43), and protection against phage predation (44). This transition is mediated by extracellular autoinducers (45, 46) which impact the activity of cytoplasmic regulators, including LuxO and HapR.

In the low-cell-density state, LuxO is phosphorylated leading to inhibition of *hapR* mRNA translation via regulatory sRNAs, whereas at high-cell-density, LuxO is dephosphorylated and inactive, resulting in HapR protein translation and the switch to high-cell-density (41, 47). Recently, a presumably laboratory acquired mutation in *luxO, luxOG319S* was discovered in some laboratory isolates of the widely used *V. cholerae* strain C6706 that locked this strain in the low-cell-density state (48). After sequence validating mutants from the 2008 library, we observed the *luxO*G319S mutation, indicating it was present in the parent strain from which the library was constructed. Thus, the 2008 ordered mutant library (32) is only partially representative of how *V. cholerae* senses and responds to its environment and a new library in a quorum sensing competent strain would greatly enhance our understanding of *V. cholerae* ecology and evolution.

To overcome this limitation of the 2008 library, we use chitin-independent natural competence to regenerate the defective ordered *V. cholerae* library in a quorum competent C6706 genomic background. To streamline the transfer process, we developed a cost effective gDNA extraction method using thymol (49). Compared to other methods, our approach is both economical ($0.64 per reaction vs $2.85 per reaction) and efficient, enabling up to 200 gDNA extractions in as little as 2.5 hours, in contrast to 25 reactions requiring >6 hours using conventional methods. Using our methods, we successfully regenerated the library with a 99% success rate, as indicated by growth on antibiotic selection growth media. Sequence validation of randomly selected mutants from the regenerated library show that the transposon insertions recombine into our recipient strain with 100% precision and transfer of the *luxOG319S* mutation never occurred. Of significance, assays performed using paired transposon mutants from each library provide evidence for density-dependent epistasis in several flagellar genes, and when grown on inosine – a metabolite secreted by gut bacteria – illustrating the utility of this new library. Lastly, we show that strains from the library generated in this work – hereafter the Grant Library – can undergo additional edits using chitin-independent natural competence. Taken together, the Grant Library represents an added contribution in our efforts toward understanding *V. cholerae* behaviors under transcriptional regulatory states that are representative of how this pathogen evolves and interfaces with its environment.

## Materials and Methods

### Bacterial matings

To generate the library recipient strain used in this study, NG001, we mated wildtype *V. cholerae* C6706 with an *E. coli* S17-λpir donor strain harboring plasmid pMMB-*tfoX*-*qstR* (generously provided by Ankur Dalia, Indiana University). In short, we mixed equal volumes of donor and recipient strains in the center of a Luria-Bertani agar plate which we incubated at 37℃ for three hours. Afterwards, we added 1 mL of LB media containing chloramphenicol (for selection of plasmid in recipient strain) and polymyxin B (for counterselection of the donor *E. coli* strain) to the plate, washed, and collected the mating spot into a borosilicate glass culture tube, which we incubated overnight at 37℃ with orbital shaking at 210 RPM.

### Media and buffers

#### Thymol cell lysis agent

We prepared thymol at a stock concentration of 350 mM by dissolving thymol crystals in dimethyl sulfoxide (DMSO). Thymol stocks were made in 15 mL which we stored at room temperature on the lab bench until they were exhausted. We observed no differences in the lytic activity of thymol during storage.

#### *gDNA* extraction *buffer*

We prepared solutions of a gDNA extraction buffer in two 500 mL volume parts. We prepared solution one by mixing 80 mL of 1M Tris (pH 8.0), 56 mL of 0.5M EDTA (pH 8.0), 46.7 g NaCl, and 3 g sodium metabisulfite, which we stirred until dissolved. We then adjusted this solution to the final volume with dH_2_O and autoclaved for 25 minutes. We prepared solution two, sodium acetate (2.5M, pH 5.2), by dissolving 102.5 g of the salt in dH_2_O, which we sterilized using filtration. Prior to each extraction, we prepared a master mixture of solution one and solution two by mixing 21 mL and 31.5 mL for every 100 samples processed, respectively.

#### Minimal media with inosine

We prepared inosine at a stock concentration of 133 mM by mixing the compound in sterile dH_2_O. We then applied gentle heating and stirred the mixture until the solution was homogenous, which we then filter sterilized. We prepared 1x M9 minimal media amended with 0.1 mM CaCl_2_, and 2 mM MgSO_4_, to which we added inosine to a 20 mM final concentration.

### Strain revival

We revived strains from the nonredundant library ((32); henceforth referred to as donor library) in microplate format to maintain parity between it and that produced in this work. The donor library was generously provided by the laboratories of Vic DiRita (plates: 2, 4-6, 8-10, 12-17, 19-33), Michigan State University, and Bonnie Bassler (plates: 1, 3, 7, 11, 18, 34), Princeton University. Briefly, microtiter plates from the donor library were slightly thawed and 10 µl from each well was inoculated into a deep-well microplate containing 550 µl of selective media (Luria-Bertani broth + Kanamycin (100 µg/ml) in each of the 96 wells. We revived the recipient library strain, NG001, in a 250 ml baffled flask containing 25 ml of selective media (Luria-Bertani broth + chloramphenicol (10 µg/ml)) to maintain plasmid *pmmB-tfoX-qstR* and 100 µg/ml isopropyl β-D-1-thiogalactopyranoside (IPTG) to induce natural competence. Unless noted otherwise, we grew all cultures overnight (18 – 24h) at 37℃ with orbital shaking at 210 RPM.

### Whole gDNA preparation

The non-redundant donor library is comprised of 35 96-well plates of *Vibrio cholerae* mutants. To extract genomic DNA from the donor library in a cost efficient and high-throughput fashion, we developed an extraction method using a modified recipe of the gDNA extraction buffer developed by Mantel and Sweigart, 2019 (accessed September, 2022, at https://www.protocols.io/view/quick-amp-dirty-dna-extraction-4r3l287zql1y/v1). In brief, we pelleted cells from overnight cultures of the donor library by centrifuging the deep-well plates at 4℃ for 20 minutes. We then removed ∼500 µl of supernatant from each well, resuspended the cell pellet in 10 µl thymol (∼35 mM final concentration), and incubated the plates at 64℃ for 15 minutes. Thereafter, we added 500 µl of gDNA buffer to each well and incubated the plates at 64℃ for an additional 45 minutes.

To clear culture supernatants of cellular debris, we transferred cell lysates from each well into an E-Z 96™ Lysate Clearance Plate (https://www.omegabiotek.com/product/e-z-96-lysate-clearance-plate/). We then used a vacuum manifold to filter the supernatant through the plate, which we collected in a second deep-well plate containing 200 µl of 100% cold isopropanol per well. We then incubated the plates for 15 minutes at room temperature to precipitate genomic DNA. Afterwards, we again used a vacuum manifold to filter the cell lysate containing our gDNA through an E-Z 96™ DNA plate (https://www.omegabiotek.com/product/e-z-96-dna-plates/?cn-reloaded=1). According to the manufacturer, “E-Z 96™ plates are silica glass fiber plates that can bind up to 50 µg of genomic DNA or 20 µg of plasmid DNA per prep.” After this initial filtration, we washed our bound DNA, in sequence, using 500 µl of 70% and 95% ice cold ethanol. We then added 100 µl of 1x TE buffer to each well and eluted genomic DNA from the DNA plates into PCR plates by centrifugation (max speed, one minute).

### Chitin-independent natural competence for gDNA transfer

After 18-24 h of overnight growth, we diluted the library recipient strain (NG001) 1:4 in 0.5x instant ocean to which we added chloramphenicol (5 µg/ml) and IPTG (100 µg/ml). We then added 200 µl of these competent cells to each well of a deep-well 96-well plate and added the gDNA extracted from the donor library to a matched recipient well (final dilution 1:3). We then incubated the deep-well plates overnight in a standing incubator at 30℃. Following the overnight incubation, we added 400 µl of selective media (Luria-Bertani broth + chloramphenicol (5 µg/mL) + kanamycin (100 µg/mL)) to each well and incubated the plates at 37℃ in a standing incubator for 48 h (Fig. 1). We determined transformation success by spot plating 3 µl of each well onto selective LB agar plates (Luria-Bertani agar + chloramphenicol (5 µg/mL) + kanamycin (100 µg/mL)), which we grew overnight at 37℃ (Fig. S1).

**Figure 1.**
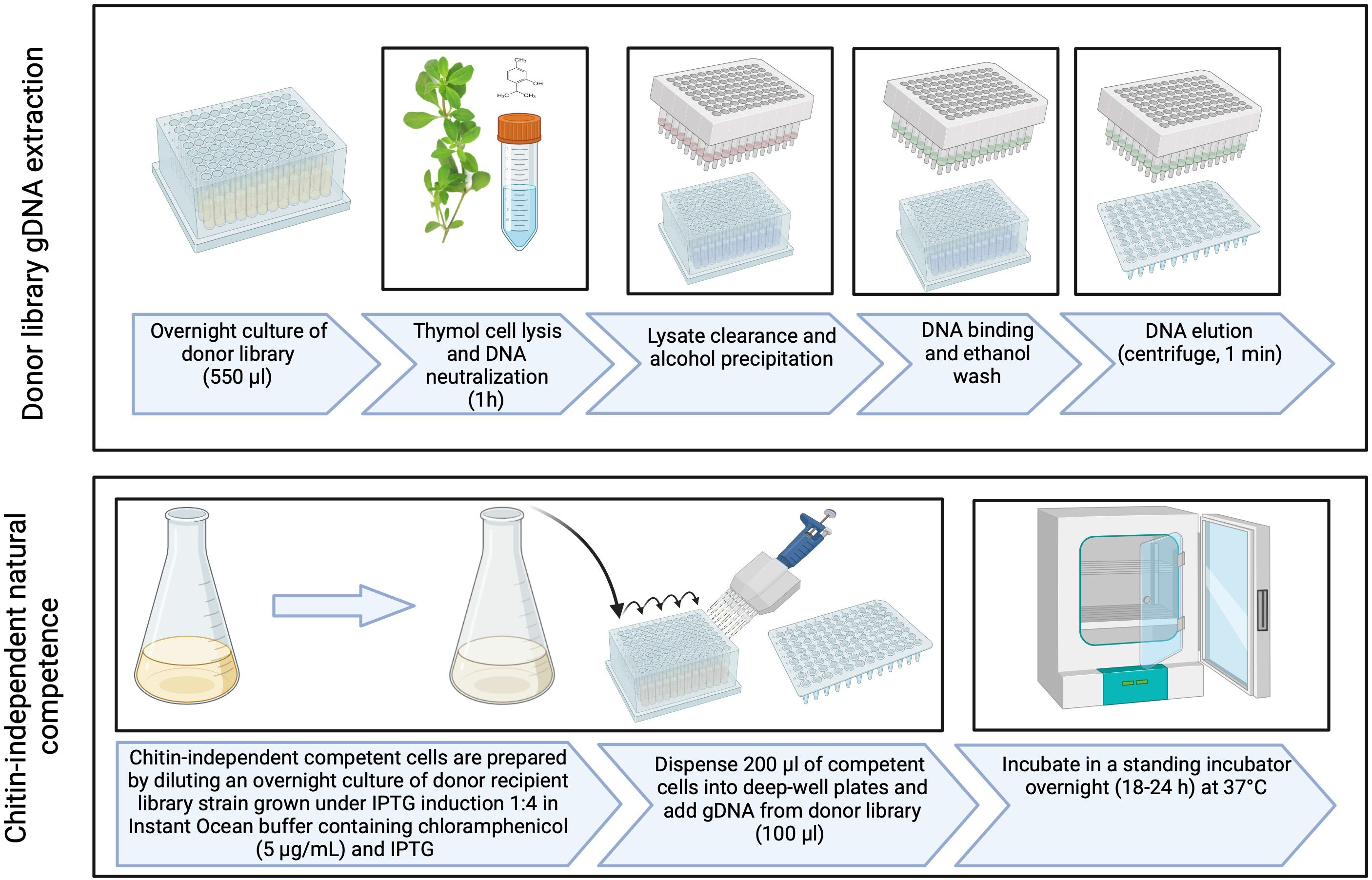
Thymol-Assisted gDNA extraction and chitin-independent natural transformation. Overnight cultures of the donor library were grown in 550 µl of selective media (LB + kanamycin). After overnight growth, the samples were centrifuged, the supernatant was aspirated, and the cell pellets were incubated for one hour at 64℃ in a novel gDNA extraction buffer containing thymol. gDNA was then isolated from the clarified supernatant using an alcohol precipitation method, followed by resuspension of the gDNA in 1x TE buffer. In parallel, a quorum-competent wildtype *V. cholerae* strain carrying the plasmid conferring natural competence (pMMB-tfoX-qstR) was grown in selective media (LB + chloramphenicol) supplemented with IPTG. The next day, competent cells were diluted 1:4 in 0.5x Instant Ocean, to which chloramphenicol and IPTG were added. Subsequently, 200 µl of cells were dispensed into the wells of a deep-well 96-well plate and co-incubated with 100 µl of our gDNA extraction at 30℃ for 18 – 24 h. For a detailed description, refer to the Materials and Methods section.

### Grant Library transformation efficiency analysis

We created a comprehensive plate indices file (Dataset S1) covering all possible wells within the donor library (P1A1 to P34D6; 3,210 wells). Using this file, we developed a script to compare it with a modified version of supplementary dataset 3 from Cameron et al., 2008 (Dataset S2), which allowed us to identify empty wells (86) and those containing mutants (3,124). Our comparative analysis also flagged duplicate entries in supplementary dataset 3 of the donor library (Dataset S3).

While constructing the Grant Library, we meticulously documented the outcomes of growth on selective media following chitin-independent natural transformation of the donor library: True Positives (3,102 wells), False Negatives (22 wells expected to grow but didn’t), True Negatives (71 blank wells that didn’t transfer), and False Positives (15 blank wells that transferred unexpectedly). These values served as inputs for a binary confusion matrix (50), providing a comprehensive assessment.

### TnFGL3 transposon insertion site identification

#### Genome Sequencing

We randomly selected 23 mutants from the Grant Library to identify the transposon insertion site. Strains were incubated overnight in 3 ml of LB + Kanamycin (100 µg/ml) and DNA was extracted from 1 ml of culture using a Wizard ® Genomic DNA Purification Kit following the manufacturer’s instructions. DNA was rehydrated overnight at 4℃ in 50 µl of TE buffer. Library preparation was performed by SeqCoast Genomics using an Illumina DNA Prep tagmentation kit and unique dual indexes. Samples were multiplexed and sequenced on the Illumina NextSeq2000 platform using a 300-cycle flow cell kit to produce 2x150 bp paired reads. Read demultiplexing, read trimming, and run analytics were performed using DRAGEN v3.10.11, an on-board analysis software on the NextSeq2000. Raw sequencing data generated during this study are available in the following public Github repository (https://github.com/NkrumahG/Grant-Library-Construction)

#### Reference *genomes*

The reference genomes for *V. cholerae* strains used in this study were downloaded from NCBI GenBank. The chromosome accession numbers for chromosomes I and II are, NZ_CP028827.1 (2,975,504 bp) and NZ_CP028828.1 (1,072,331 bp) for strain N16961, and NZ_CP046844 (3,019,938 bp) and NZ_CP046845 (1,070,359 bp) for strain C6706, respectively. The sequence map for plasmid pSC189 was downloaded from addgene (https://www.addgene.org/browse/sequence/36279/), and the sequence for pMMB-*tfoX*-*qstR* was provided by Ankur Dalia (Indiana University).

#### Mutation *calling*

We used consensus mode in *breseq* ((51), version 0.37) to align the paired-end reads for each of the sequenced mutants to the reference genomes. The computational resources used to perform *breseq* were provided by the Institute of Cyber-Enabled Research at Michigan State University.

#### *Ortholog* identification

We used Mauve ((52), snapshot _2015-02-25) to identify orthologs in whole genome sequences of C6706 and N16961. Each pair of chromosomes were aligned using the progressive Mauve method with default settings. In the Supplemental Information, we have included lists of all orthologous gene pairs in *V. cholerae* N16961 and C6706 (Dataset 4) and the identity of genes unique to N16961 (Dataset S5) and C6706 (Dataset S6). Additionally, we used Geneious to extract, align, and visually display (Supplementary Information, Fig. S2 and S3) orthologous gene pairs for the transposon mutants sequenced in this study.

### LuxO gain of function mutation identification

Using supplementary dataset 3 provided for the donor library (32), we identified all potential transposon mutants located within a 50 kb range, both upstream and downstream of *luxO*. Following this selection, we cultured these 92 mutants in selective media (LB + Kanamycin, 100 µg/mL) for 24 hours under standard conditions.

After the incubation, we extracted gDNA using the thymol-based gDNA extraction method we developed to construct the Grant Library. The extracted genomic DNA was then diluted 1:2, which we used as template for PCR amplification of the *luxO* gene. The *luxO* gene was amplified with Q5 polymerase using the following primers: forward primer 5’ GGCTATGCAACATAATCAATCTTG-3’ and reverse primer 5’-GCTTTGGTTGATCCATTCTCTCAT-3’. PCR conditions included an initial denaturation at 98℃ for 2 minutes, followed by 29 cycles of denaturing at 98℃ for 30 seconds, annealing at 63℃ for 30 seconds, and extension at 72℃ for 1 minute. A final extension step was performed at 72℃ for 5 minutes.

Following the PCR amplification, we validated amplification of *luxO* using gel electrophoresis. We then used the Qiaquick PCR purification system to purify our *luxO* PCR product. Sanger sequencing of the purified *luxO* allele was performed using the reverse primer 5’-TGCGATAGATGGTTGACGGG-3’ by the Roy J. Carver Biotechnology Center at the University of Illinois at Urbana-Champaign. We imported the Sanger Sequence read data into Geneious (53) and aligned it to *luxO* (*V. cholerae* chromosome I, C6706), allowing comprehensive analysis and visualization of our sequenced mutants, with special attention to amino acid position 319, the site of the *luxO* gain of function mutation *luxOG319S* (48).

### Motility Assay

We screened mutants with presumptive TnFLG3 transposon insertions in known motility genes. Strains were grown in a deep-welled 96-well plate overnight at 37℃ in selective media (LB + Kanamycin) and on the following day, we used toothpicks to manually stab each strain into motility plates containing LB and 0.35% agar. Plates were incubated at 37℃ for 24 h, and the motility zones were recorded with a gel imager. The area of the motility zones was measured computationally using the Hough Circle Transform package from the UCB vision plugin in Fiji (54).

### Inosine Growth Assay

We randomly selected 5-paired 96-well plates from each of the Grant and Cameron et al. transposon mutant libraries and incubated them overnight in LB plus kanamycin (100 µg/mL). We also revived the parent strains used to construct each library in the same fashion. On the following day, we recorded mutants that didn’t grow overnight and then back diluted each well from the plates 1:100 into M9 minimal media supplemented with 20 mM inosine. We incubated all plates with linear shaking at 37°C. We monitored growth of the parent strains at an OD_600_ for 10 hours, with measurements taken at 5-minute intervals. For the paired library mutants, we measured the 10 h endpoint optical density.

### Plasmid Curing Assay

We inoculated three cultures of NG001 from frozen glycerol stocks into LB media supplemented with chloramphenicol (10 µg/ml) and incubated them overnight. On the following day, we back diluted each culture 1000-fold into either LB, LB media containing chloramphenicol (10 µg/ml), or LB containing IPTG (100 µM). For the duration of the experiment, we passaged each culture in the same media. In parallel, we serially diluted stationary phase cultures in 0.5x Instant Ocean and spot plated 3.0 µl from each dilution onto LB or LB agar containing chloramphenicol. The percent of population with the *pMMB-tfoX-qstR* plasmid (plasmid carriage) was calculated as a ratio of the dilutions where colonies grew on LB with chloramphenicol relative to growth on LB alone.

### Data accessibility

All data generated, and analysis scripts used in this work are available in the GitHub repository at https://github.com/NkrumahG/Grant-Library-Construction.

### Results and Discussion

#### Chitin-independent transfer of transposons from donor library strains into a quorum-competent *V. cholerae* genetic background is remarkably efficient

We hypothesized that the transposon insertions within the genomes of the quorum-incompetent *V. cholerae* strain generated in Cameron et. al., 2008 could be transferred to a quorum-competent wildtype *V. cholerae* strain using natural competence. To test this, we extracted gDNA from each mutant in donor library using an in-house method involving thymol lysis and alcohol precipitation. Subsequently, we co-incubated this gDNA with a wildtype *V. cholerae* strain (C6706) carrying a plasmid that expresses the natural competence master regulator, TfoX, upon induction with IPTG (Fig. 1, Materials and Methods).

The donor library consists of 34 plates arrayed from P1A1 to P34D6, for a total 3,210 possible wells containing transposon mutants. Upon examination of supplementary data table 3 for the donor library, we discovered that 86 wells were not listed, and some of the mutants were recorded multiple times. With these considerations, we expected growth for 3,124 unique transposon mutants after conducting our natural transformation protocol.

To verify the presence of transposon insertions in the Grant library mutants, we spot-plated each mutant on LB medium supplemented with kanamycin. Of the 3,124 unique transposon mutants we expected to grow on the plates, 3,102 mutants (99.2%) exhibited growth (Fig. S1). None of the 22 missing mutants, which were expected to grow, showed growth in selective media during overnight revival of donor library freezer stock. This suggests that these mutants might no longer viable perhaps due to low culture density upon freezing or death resulting from repeated freeze-thaw cycling during long-term storage of the donor library. In the cases where we anticipated no growth, 71/86 (82.0%) did not grow on selective media.

To evaluate the performance of our library transfer approach, we utilized a confusion matrix (Fig. 2). The results of this analysis indicate that our approach was highly accurate and precise, with a high recall (F1 score = 0.9941) and a low False Discovery Rate (0.0070). These results unequivocally demonstrate the remarkable efficiency of using chitin-independent natural competence to transfer the transposons into a quorum-competent strain. The transference process was made possible through our in-house gDNA extraction method, involving thymol cell lysis and alcohol precipitation, affirming the suitability of this gDNA preparation for downstream applications. Indeed, this suitability was confirmed when we employed thymol-extracted DNA as a template in PCR, with no observed inhibitions.

**Figure 2.**
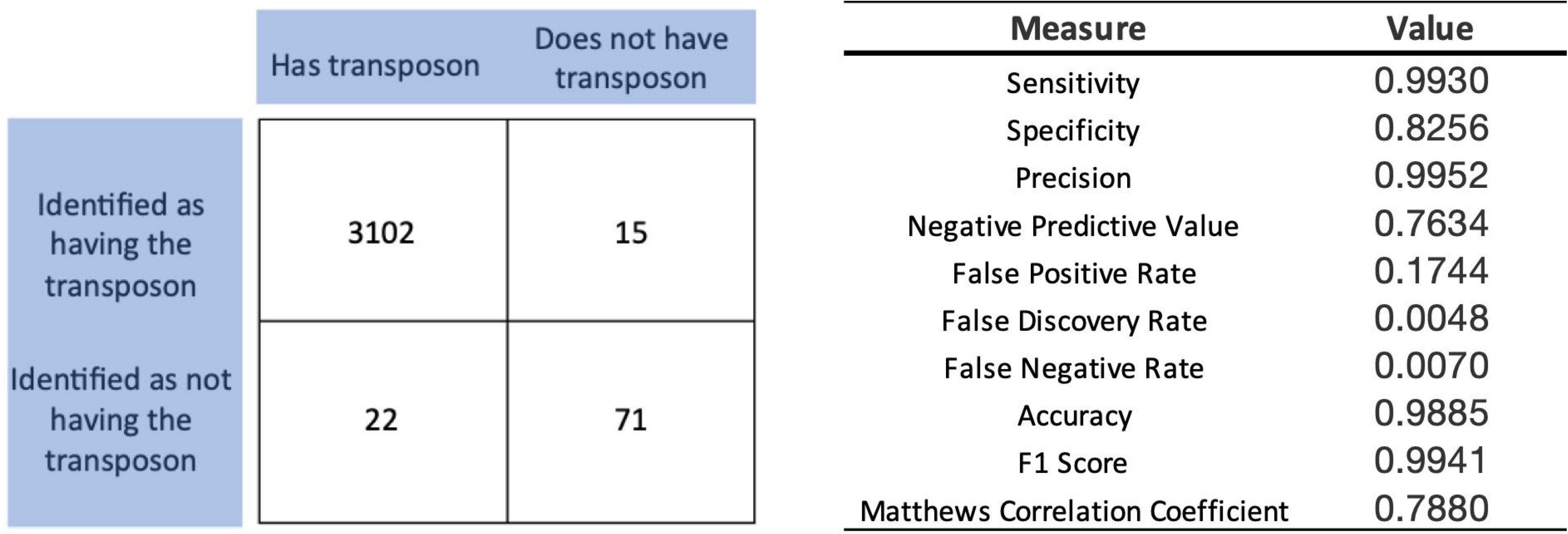
Analysis of Library Transfer. The outcomes of growth on selective media for each mutant were recorded after chitin-independent natural transformation of the donor library. These values were utilized as inputs to construct a 2x2 confusion matrix, categorizing the results as follows: True Positives (3,102 wells), False Negatives (22 wells), True Negatives (71 wells), and False Positives (15 wells). The accompanying table presents the calculated measures, the values of which were computed using standard derivations.

#### Transposons from the donor library homologously recombine into the recipient genome with 100% precision

The frequency of genome edits by homologous recombination in *Vibrio spp.* is highest when there are at least 2 kB arms of homology flanking the genomic sequence being altered (25). Given this requirement, the observation that the recipient strains grew after incubation on selective media (Fig. S1) was suggestive that transposons correctly integrated into the homologous site of the recipient’s genome. To validate our supposition, we randomly selected 23 mutants, isolated gDNA, and sequenced whole genomes. Subsequently, we aligned the sequenced reads to four reference sequences using breseq (51). The reference sequences included *V. cholerae* chromosomes I and II, and the plasmid conferring natural competence. A fourth reference sequence (pSC189) containing sequences associated with the transposon was also included as a “genomic lure” to capture reads that mapped to the transposon. When examining the breseq output, we specifically looked for instances where new junctions were created between chromosomes I or II (depending on which chromosome the gene targeted for homologous recombination was located) and pSC189. Such a junction would indicate a transposon integration event. Fig. 3 is an example breseq output from one such alignment, which reports the transposon insertion site in chromosome II, including the basepair position it is located.

**Figure 3.**
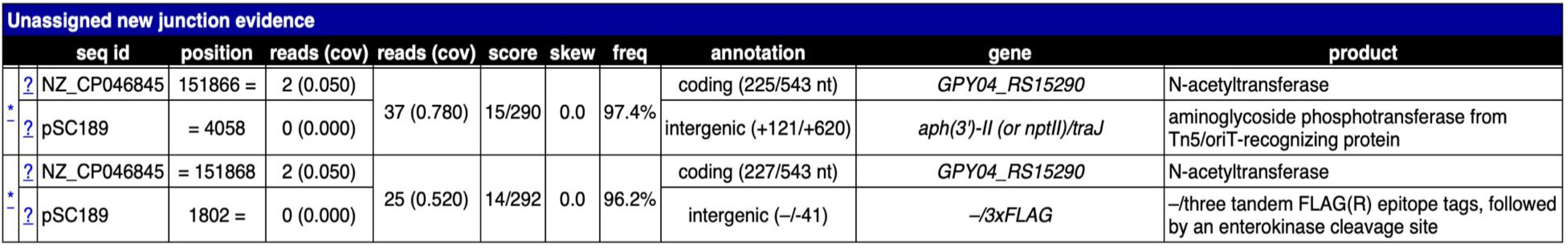
Representative sample of breseq output showing new junction evidence supporting transposon insertion. The figure shows a portion of the summary.html file generated for a breseq analysis run. In this example, a new junction has been called between chromosome II (NZ_CP046845) of *V. cholerae* strain C6706 and the plasmid (pSC189) containing representative sequences of the transposon. Insertions have left and right boundaries, where the chromosome and sequence meet. The boundaries are marked with an asterisk. The annotation column shows the position of the transposon within the gene (GPY04_RS15290), which we converted to a value called “percent into open reading frame.” In this example, that value is 41.43% ((225/543) * 100)). This value was calculated for all strains sequenced in this study.

The parent strain of both the donor and recipient libraries is *V. cholerae* El Tor C6706. However, when the donor library was generated in 2008, transposon insertion sites were mapped to reference genomes of *V. cholerae* El Tor strain N16961 because it was the only *V. cholerae* that had been sequenced for *Vibrio spp.* at that time (55). In this work, we mapped transposon insertion sites to reference genomes of strain C6706. To identify whether transposons insertions were in the correct site of our recipient strains, we performed whole genomic comparative analysis of *V. cholerae* N16961 and C6706. When we performed this analysis, we found chromosomes I and II differ in size between strains and are not syntenic (Fig. 4). Out of the 2,649 genes annotated in our reference genomes for chromosome I, 2,558 (96.6%) of genes are shared between N16961 and C6706. Additionally, 21 genes (0.79%) are unique to N16961 and 70 (2.64%) to C6706. On chromosome II, 1,052 out of 1,064 annotated genes (98.9%) are shared between strains, where 10 (0.94%) and 2 (0.19%) of genes are unique to N16961 and C6706, respectively. Importantly, this analysis provided the genomic position for each ortholog of N16961 genes present in C6706 (Dataset S1).

**Figure 4.**
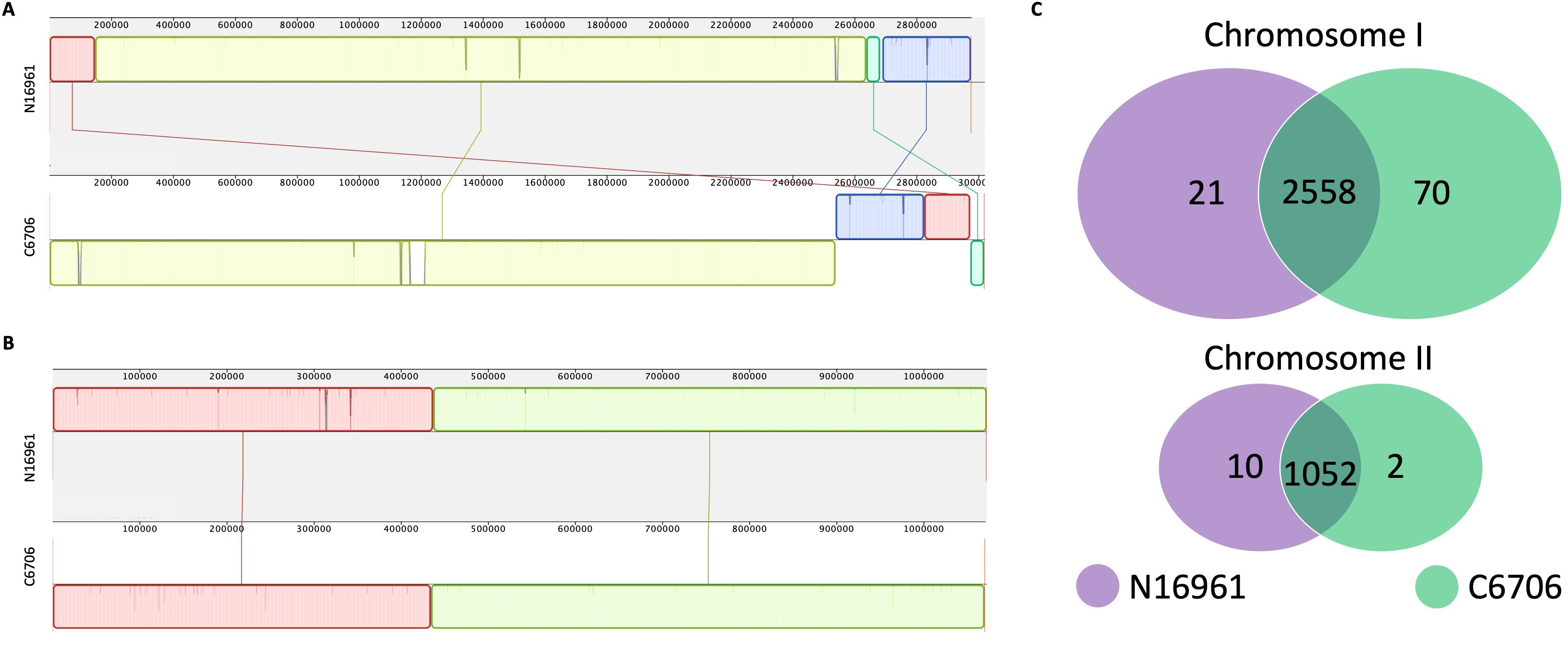
Genomic comparative analysis of *V. cholerae* N16961 and C6706 for chromosome I (A) and chromosome II (B). Genbank files for whole-genome sequences of chromosomes I and II of both strains were downloaded from NCBI. The analysis involved a comparative examination of genetic variations and conservation patterns between the two strains. *V. cholerae* strain N16961 is shown with a gray background and C6706 a white one. Genome segments are shown as colored colinear blocks centered around the “positive” and “negative” strands of each chromosome. Lines connecting the segments show their orthologous alignments between the strains, including inversions. Panel C is a Venn diagram summarizing the relationship of genes for each strain and chromosome. The data underlying the illustration in panel C, including the genomic position of orthologous gene pairs, is provided in the Supplementary Information (Datasets S2–S4).

We used the orthologous sites to analyze each transposon mutant we sequenced (Dataset S1, Fig. S2 and S3). Using the breseq annotation data, we calculated the position of the transposon within each gene as a percent of basepairs into the ORF of C6706. These position values were previously reported for each strain in the donor library strain N16961. To assess the relationship, we performed a linear regression of our sequenced C6706 insertions with the previously described N16961 from the donor library, expecting a perfect linear correlation. Although 19/23 mutants had an exact match, some of our values diverged from the line of best fit (Fig. 5), although there was a strong statistical correlation between the variables (Kendall’s coefficient τ = 0.752, *n* = 23, and *P* < 0.0001). Upon further investigation of this anomaly, we discovered two categories of false negatives. In category one (Fig. S3, Panels A and B), we observed two cases where genes were annotated in N16961 but not in C6706. Consequently, these data points lacked a calculated value for the transposon insertion site in C6706 and thus fell on the y-axis. In the second category, two false negatives were attributed to a difference in the size of the gene orthologs between N16961 and C6706 (Fig. S3, Panels C and D). This discrepancy resulted in a shift of the relative position reported for the transposon. After accounting for these deviations and correcting the data accordingly, we confirmed that the transposon inserted into the cognate position of the Grant Library mutants we sequenced with 100% precision.

**Figure 5.**
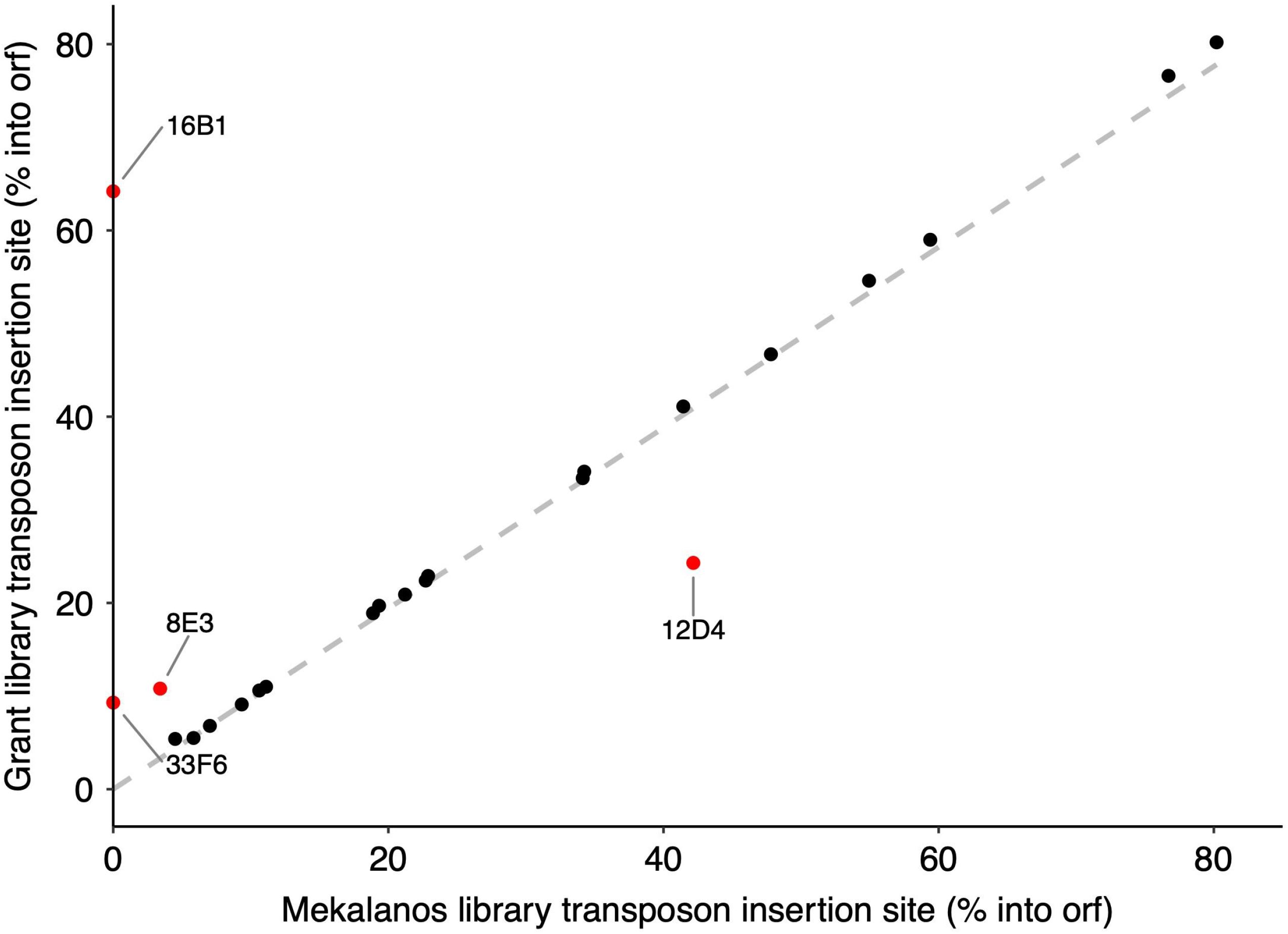
Correlation between transposon insertion sites in *V. cholerae* N16961 and C6706. Each point represents the transposon insertion site, calculated as percent into open reading frame. The values for strain N16961 were reported in supplementary dataset 3 of Cameron et al. (2008), and in this work, they were calculated using the data provided in the annotation column of the breseq output (Fig. 3). The correlation analysis yields a strong positive correlation with Kendall’s coefficient τ = 0.752 (*n* = 23, and *P* << 0.0001), indicating a significant relationship between the insertion sites of the two strains.

#### Co-transformation of the *luxOG319S* gain of function mutation does not occur

Since we employed natural competence to transfer the transposon mutations, there is a possibility that the *luxOG319S* mutation from the donor library might have been co-transformed along with the transposon into the Grant library. However, unlike the strong selection pressure for the transposon mutation, selection for the *luxO* mutation would be indirect and weaker, depending on whether the *luxO* mutation provides a fitness advantage to cells allowing them to outcompete wildtype cells during outgrowth in selective media.

In the 23 genome sequences analyzed above, none had the *luxOG319S* mutation. However, we did consider that the frequency of transfer of *luxO* mutation could be elevated in genes for which the transposon is within close genomic proximity to *luxO.* To test this, we isolated gDNA from all 94 mutants within a 50kb window up- and downstream of *luxO* (Dataset S7). We then PCR amplified *luxO* and performed Sanger sequencing. In no instance did the *luxOG319S* gain of function mutation transfer from the donor to the recipient library parent strain (Dataset S8). This suggests that proximity of the transposon mutations to *luxO* does not influence transfer of the gain-of-function mutation, however we do recommend sequencing *luxO* as a best practice when drawing comparisons between donor and recipient library strains.

#### Natural competence in Grant library strains is IPTG inducible

When generating the Grant library, we supplemented the outgrowth media with chloramphenicol and kanamycin to select for maintenance of the plasmid conferring IPTG-induced natural competence, pMMB-*tfoX*-*qstR*, and the TnFLG3 transposon, respectively. We reasoned that a library established with strains harboring this plasmid would be advantageous to the research community as it would facilitate additional genome edits using chitin-independent natural competence and homologous recombination upon induction with IPTG. In proof of concept for this idea, we randomly selected 10 mutants from the Grant library and performed a knock-in experiment of a trimethoprim (tm) resistance gene. In short, we used thymol to isolate whole gDNA from a tm resistant strain and added 16.2 µg of whole gDNA to our competent cells, which we prepared following our protocol described in Materials and Methods.

After incubating our competent cells with gDNA overnight, we used spot plates to assess antibiotic phenotypes. All the randomly selected library mutants grew on LB plates supplemented with trimethoprim between dilution factors of 10^-3^ – 10^-8^ (Fig. 6). When also plated on growth media supplemented with kanamycin, there was parity in the growth phenotypes observed on tm plates for all but one mutant (strain 8C5). When we examined the images of the kanamycin supplemented spot plates when the library was constructed (Fig. S1), we found mutant 8C5 didn’t grow there either. We posit that the spurious growth of strain 8C5 likely reflects a frozen subpopulation of our library recipient strain that was able to withstand kanamycin during outgrowth or contamination introduced during the spot-plate assay. Taken together, all the library mutants we tested could undergo additional genome edits with overnight induction of natural competence.

**Figure 6.**
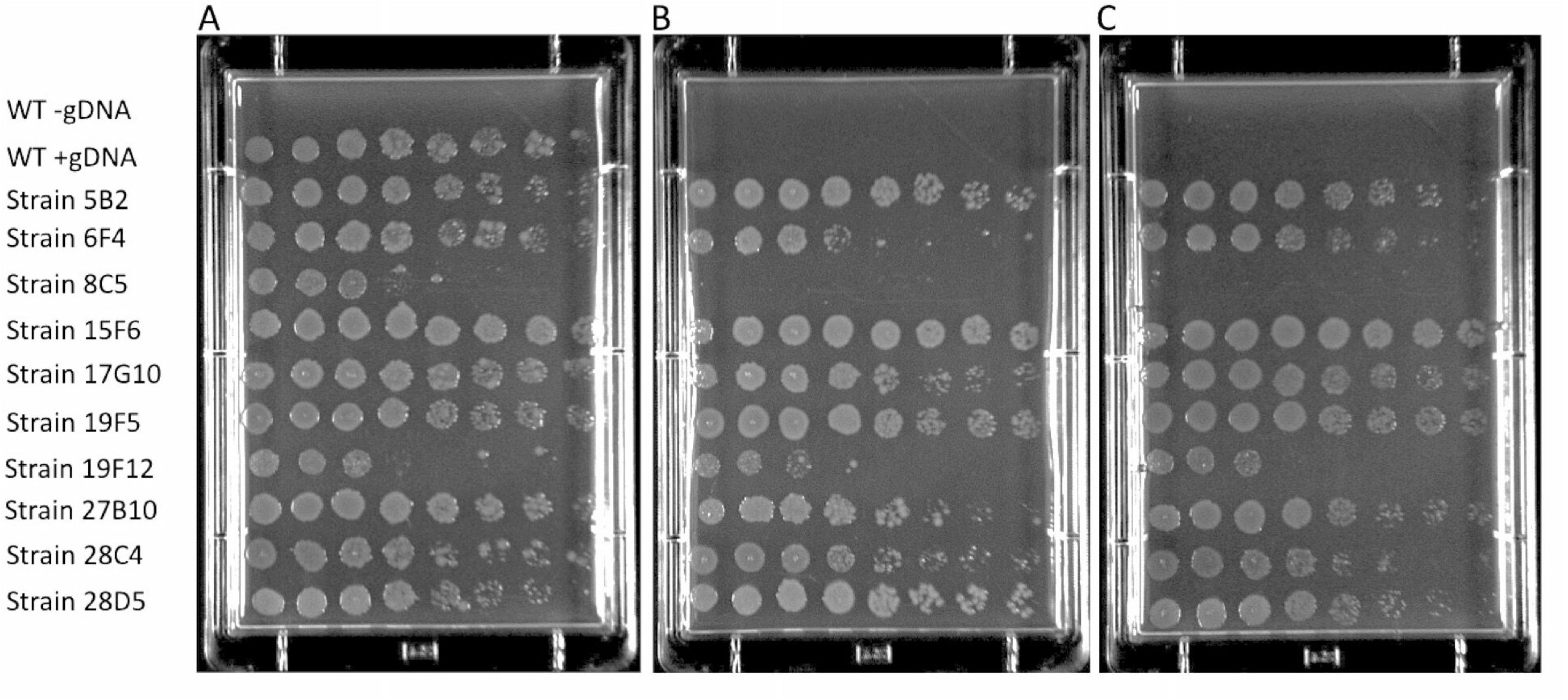
Chromosomal integration of a trimethoprim resistant gene in select mutants from the Grant library using IPTG-induced natural competence. We induced natural competence in wildtype *V. cholerae* strain C6706 carrying plasmid pMMB-*tfoX*-*qstR* (NG001) and 10 randomly selected mutants from the Grant library according to procedure described in Materials and Methods. These strains were then co-incubated with gDNA extracted from a strain carrying a trimethoprim resistant cassette and on the following day spotted on antibiotic selection media. All the strains carrying the plasmid grew on plates supplemented with trimethoprim (Panel A). On the plate with kanamycin (Panel B) or kanamycin + trimethoprim (Panel C), all strains grew except strain 8C5, indicating that this strain does not have a transposon insertion.

#### Plasmid conferring natural competence is readily cured from Grant library strains with IPTG induction

Selecting for Grant library transposon insertion mutants retaining the pMMB-*tfoX-qstR* plasmid allows for additional genomic edits to be constructed in each strain using chitin-independent natural competence. However, plasmid carriage might impose fitness costs that can vary between mutants due to genetic conflicts with each disrupted gene or complicate efforts to introduce alternative plasmids. Taking this into account, we sought to establish whether we could cure Grant library strains of the plasmid. To this end, we serially passaged the recipient strain (NG001) carrying the plasmid in 1) LB media, 2) LB supplemented with chloramphenicol and not IPTG, or 3) LB without chloramphenicol but supplemented with IPTG. We hypothesized that amplifying plasmid carriage cost by inducing its expression in the absence of antibiotic selection (treatment three) would lead to rapid curing.

In agreement with our hypothesis, the number of cells carrying the plasmid decreased by ≈ 94% after 120 h of serial passage in LB media supplemented with IPTG alone (Fig. 7). In contrast, plasmid carriage in cells passaged in LB alone or in LB containing chloramphenicol decreased by ≈ 39% and ≈ 11%, respectively. These data show that the pMMB-*tfox-qstR* plasmid can be readily cured from Grant library strains.

**Figure 7.**
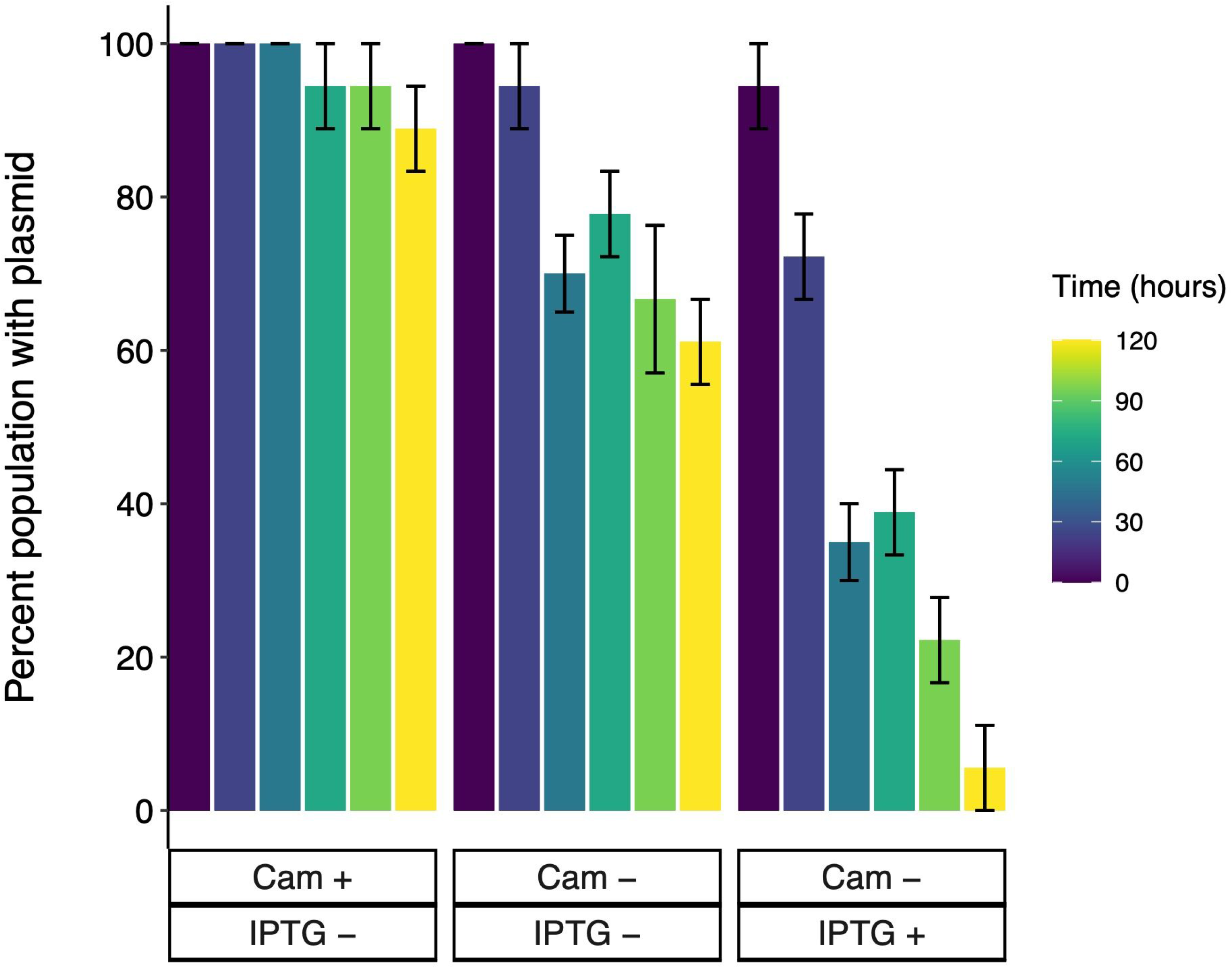
Effects of media composition on plasmid loss in *V. cholerae*. We serially propagated cultures of the Grant Library parent strain in LB media with or without antibiotics or IPTG induction as indicated in the figure. The percent of population with the plasmid conferring natural competence was calculated as a ratio of the dilutions where colonies grew on selective media (LB + chloramphenicol (Cam) relative to non-selective media (LB). Error bars are the 95 % confidence intervals on a sample size of *n* = 3 for each treatment.

#### Differential motility responses of recipient library mutants reveal unappreciated epistasis exists in donor library

1. *V. cholerae* exhibits high motility by virtue of a single polar flagellum, a characteristic that contributes significantly to its pathogenesis. *V. cholerae* isolated from active infections display increased motility compared to lab-grown strains (56) and demonstrated a heightened ability to colonize hosts (57). Moreover, motility allows *V. cholerae* to survive in its aquatic environment, enabling it to swim freely as planktonic cells or form biofilms in response to environmental stress. Furthermore, previous work has demonstrated the involvement of at least 40 genes in flagellum biosynthesis and motility, classified into temporally distinct classes (I through IV) (58–60). Notably, some of these genes are known to trigger an immune response (56).

Motility is an easily screened phenotype, and the functional consequences of null flagellar gene mutants are well established. Therefore, Cameron et al., 2008 (32), employed motility assays as one of the validation metrics for their constructed library. In this context, we also measured motility of the Grant library null flagellar gene mutants. By performing the assay using paired mutants from the donor and recipient libraries, we reasoned that we would foremost independently confirm the motility phenotypes reported by Cameron et. al., 2008. Furthermore, previous studies have indicated that genes in classes III and IV of the flagellum biosynthesis hierarchy exhibit cell-density dependent phenotypes (61). Thus, the motility assay would serve to provide supporting evidence for or against our hypothesis that quorum sensing epistasis is a missing feature of the donor library.

Out of the 33 genes examined, we successfully replicated the motility phenotype reported for the donor library’s nonmotile mutants in all but one strain (*fliR*) (Fig. 8). Additionally, we observed that both libraries exhibited nonmotile behavior for *rpoN* (class I) and *flaA,* (class III) that are essential for expression of the flagellar biosynthesis genes (Fig. 8). Furthermore, motility was also abolished for the σ-54 dependent transcriptional regulator transposon mutants of *flrA* (class I) in both libraries, consistent with previous observations (62). When comparing motility within classes, paired null mutants from the Cameron and Grant libraries displayed similar movement on the plates for all classes, except for class III, where a statistically significant difference was observed (Table 1, two-tailed *t*-test: *P* = 0.012). Additionally, two mutant pairs (*flgM* and *flgN*) within class IV biosynthesis genes exhibited opposite motility phenotypes.

**Figure 8.**
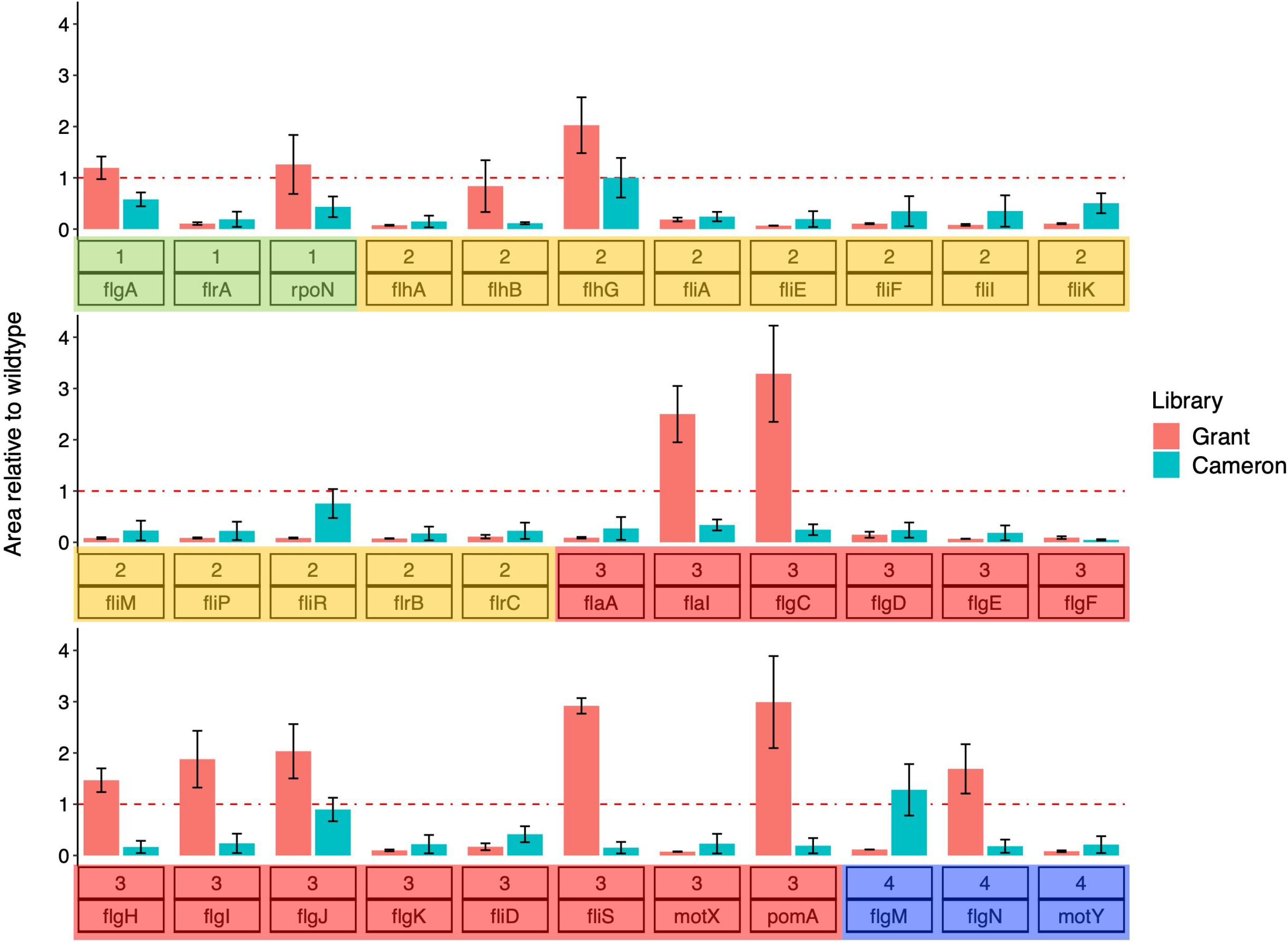
Motility behavior of flagellar mutants from the Grant and Cameron et al. (2008) libraries. Paired mutants with transposons in flagellar genes from both libraries were screened for motility in nutrient-rich plates containing 0.35% agar. The plates were imaged, and the area moved by each mutant from an inoculum site was calculated using the procedure described in the Materials and Methods. These areas were then normalized to a value relative to the wild-type strains for each library from 3 replicate assays in all but four cases (*flgM*, *flhB, fliR*, and *flgF*), which had two replicates each. The dashed red line is the normalization for the wildtype strains. Colored boxes highlight the class each gene belongs to in the flagellar biosynthesis gene cluster hierarchy, and red stars highlight the expected non-motile phenotype for transposon insertions that lie within genes encoding sigma factors required to transcribe downstream flagellar biosynthesis genes.

While these opposing effects were observed within the same class, the net effect of class IV genes canceled each other out in the statistical analysis, so this class comparison was insignificant. Taken together, our results suggest quorum sensing epistasis with motility genes results in the different motility outcomes as the donor library is locked in a low-cell density genomic background.

#### Growth on inosine is quorum dependent

Inosine is an essential nucleoside that plays an important role in purine biosynthesis and gene translation (63). Recent studies have demonstrated that various species of gut bacteria, including *Lactobacillus* (64) and *Bifidobacterium*, secrete inosine in significant quantities during growth (65). This production not only contributes to systemic immunomodulatory and protective functions (66) but also enhances mucosal barrier functions within the gut (65). Additionally, inosine has been found to inhibit the expansion of pathogenic *Enterobacteriaceae* through PPARγ activation, further highlighting its importance in preventing gut dysbiosis (67).

In an unrelated screen analyzing for quorum-dependent growth effects using Phenotype Microarrays for Microorganisms (Biolog), we observed wild-type C6706 and a locked high-cell density (Δ*luxO*) mutant exhibited enhanced growth when grown on inosine as the sole carbon source compared to a locked low-cell density mutant (Δ*hapR*). As that the *luxOG319S* gain-of-function mutant locks the Cameron library into the low-cell density state, we reasoned screening of mutants in the Grant library for growth on inosine could yield previously unappreciated epistatic effects between quorum sensing and central metabolism. Given *V. cholerae’s* manifestation of disease as an enteric pathogen – where inosine production is high – we sought to investigate whether these differences extended to other mutants from the recipient and donor transposon libraries. This exploration may shed light on the pathogenesis of *V. cholerae* and its interaction with inosine metabolism. Moreover, any significant differences between the transposon mutants would underscore the importance of using a library that doesn’t mask phenotypic states relevant to *V*. cholerae’s ecology and evolution.

We began our investigation by Sanger sequencing the *lux*O locus of the parent strains for the donor and recipient libraries. The sequence chromatograms (Fig. 8*A*) validated the glycine-to-serine gain-of-function mutation at amino acid position 319 in the donor library parent. We then grew replicate cultures of the parent strains in M9 minimal media supplemented with 20 mM inosine (Dataset S9). There was no significant difference in the growth rates of the parent strains. However, the recipient library parent strain bearing the ancestral *luxO* allele grew to an endpoint optical density that was on average ≈ 23% higher than the donor library parent strain (Fig. 8*B*) and the relationship was statistically significant (Two-Sample unpaired *t* = 29.34, *df* = 16, and *P* < 0.0001).

We then randomly selected and revived five-paired plates from the donor and recipient libraries and grew them in inosine for 24 h (Dataset S10). In agreement with the experiments performed with the parent strains, the Grant library transposon mutants grew to an endpoint optical density that was 35% higher than the Cameron et al. library (Fig. 8C) and the relationship was statistically significant (Two-Sample unpaired *t* = -20.34, *df* = 904, and *P* < 0.0001).

Reaction norms describe the phenotypic expression of a single genotype across an environmental gradient (68). Given our experimental design and the isogenicity of the donor and recipient strains, we adopted this method to visualize deviations between paired library mutants grown on inosine (Fig. 9C, grey lines). We focused our attention on mutants whose growth on inosine were above and below two standard deviations (top and bottom 2.5% of the data, respectively) away from the mean for each library (Fig. 10).

**Figure 9.**
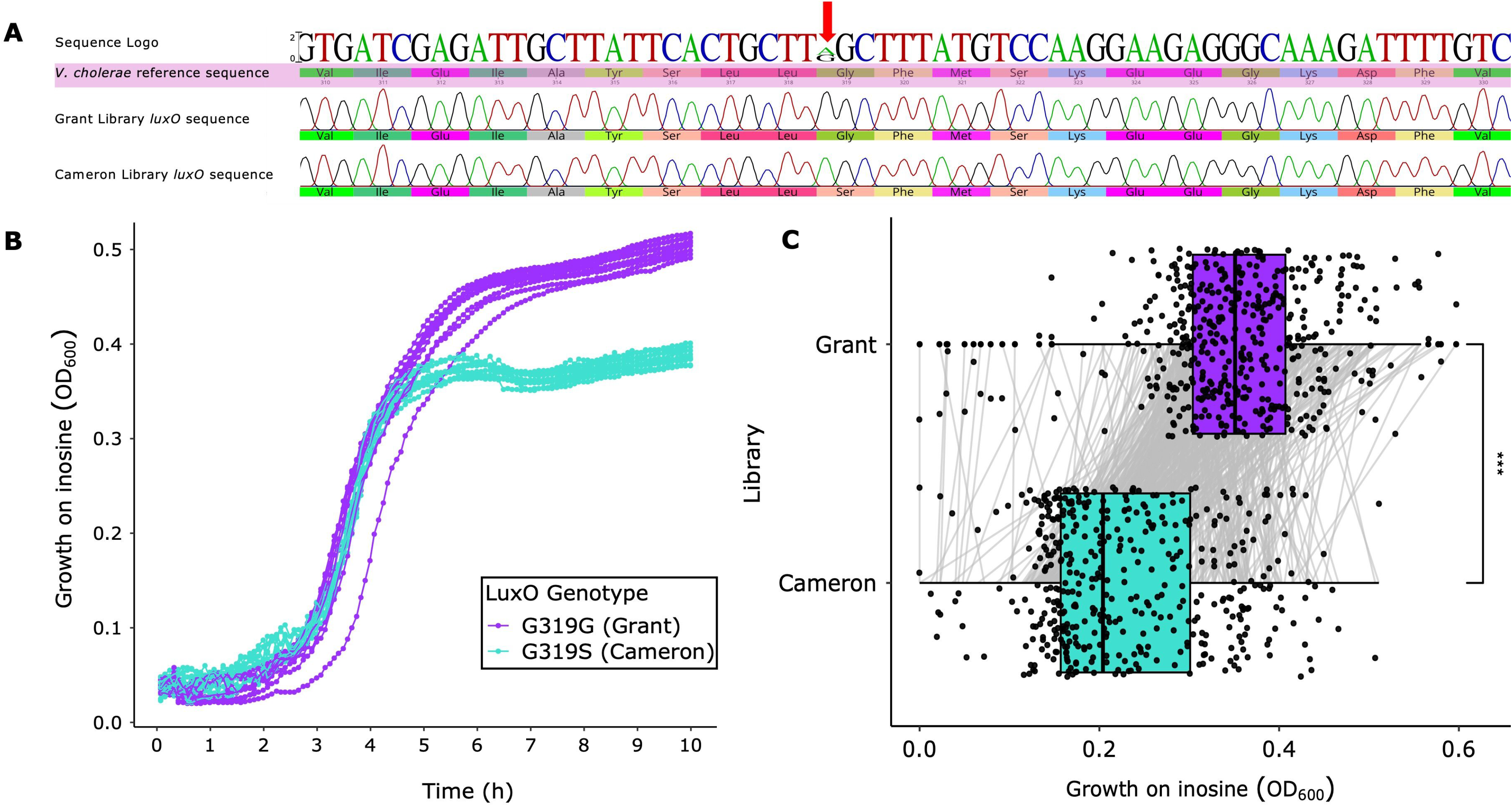
Genotypic and phenotypic characterization of Grant and Cameron library *V. cholerae* strains on inosine. (A) *luxO* sequence chromatogram for *V. cholerae* C6706 parent strains of Grant (recipient) and Cameron (donor) transposon libraries. Red arrow shows location of gain-of-function mutation resulting in a Glycine-to-Serine transition at amino acid position 319 in the Cameron library. (B) Growth of parent strains on inosine. Quorum-competent Grant library strain grows to a higher OD (Two-Sample unpaired *t* = 29.34, *df* = 16, and *P* << 0.0001). (C) Boxplots of end-point optical density values for paired mutants from Grant and Cameron libraries grown on inosine. Each point represents a single mutant and the connecting lines the reaction norm for each mutant with respect to the library sampled from. Like the parent strains (B), Grant Library mutants grows to a higher OD than the Cameron et al. Library strains (Two-Sample unpaired *t* = -20.34, *df* = 904, and *P* << 0.0001).

**Figure 10.**
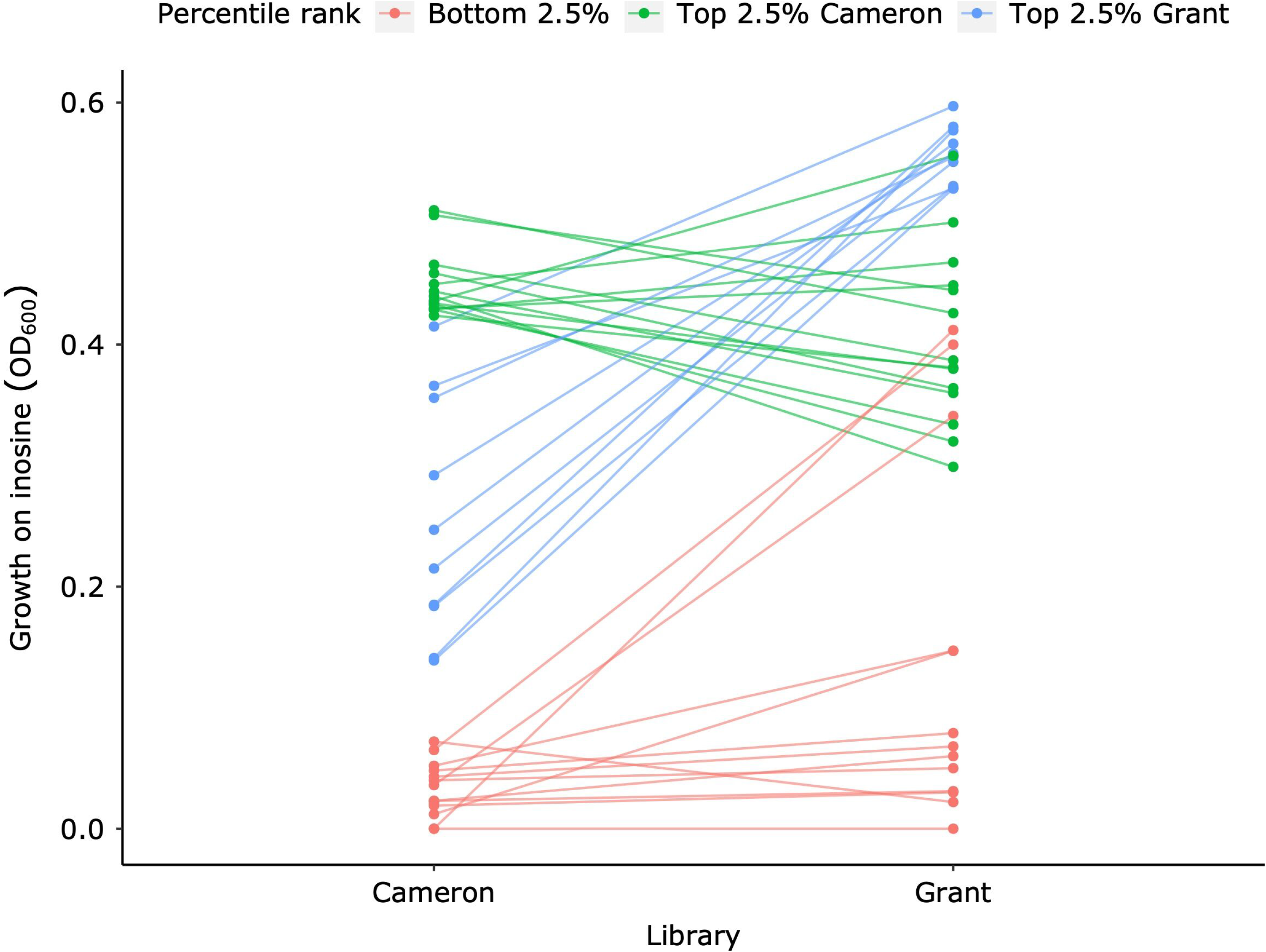
Reaction norm for paired transposon mutants grown on inosine. Each point on the graph represents the 10-hour endpoint optical density for mutants from the Cameron and Grant libraries grown on inosine, with lines connecting paired mutants. We identified outliers by focusing on the top and bottom 2.5% of the data, calculated using the mean and standard deviation for each library. Relationships are plotted for points that deviated two standard deviations from each mean.

The data shows strong evidence of quorum dependent epistasis. When we looked at the bottom 2.5% of all the data, seven of the thirteen transposon mutants were shared between libraries and each failed to grow on inosine (Tables S1 – S2). The functional roles for the associated genes included amino acid biosynthesis, energy metabolism, and regulatory functions. That these mutants failed to grow independent of the *luxO* genetic background likely reflects the essentiality of these genes when inosine is used as the sole carbon source. For the remaining six genes, five grew better on inosine in the Grant Library genomic background and one marginally less. Interestingly, when examining the top 2.5% of the data for Cameron or Grant Library paired mutants, there was strong anticorrelation. That is, where growth was high for a mutant in one library, its growth was lower in the opposite. In stark contrast to the bottom 2.5% of genes, only one transposon mutant (gene disruption of *fusA-2*), was shared within the top 2.5% of each dataset, and there was more variation in the functional roles of all called mutants (Tables S3 – S4). We hypothesize that the non-linearity in the top 2.5% is due to differences in the transcription of genes at low and high cell densities.

Collectively, our inosine findings show that ecologically relevant phenotypes can be masked for *V. cholerae* in quorum dependent contexts and are supported by several other research groups who as well have observed nuanced phenotypes in the *luxOG319S* genetic background (48).

## Conclusion

Bacteria employ quorum sensing (QS) to regulate gene expression based on population density. In the case of *V. cholerae*, the etiological agent of cholera, QS controls various phenotypes as the population transitions between low and high cell density. Researchers previously constructed an ordered mutant library, systematically disrupting every non-essential gene in *V. cholerae* with a transposon insertion. However, unbeknownst to them, this library was created in a strain with a mutation that rendered the mutants incapable of transitioning between low and high cell density.

In this study, we successfully transferred transposon insertions from these *V. cholerae* non-redundant ordered mutants into a wildtype genetic background using chitin-independent natural transformation. The resulting Grant Library comprises 3,102 mutants, covering approximately 79.8% of the ORFs annotated in *V. cholerae*.

In addition to encoding a functional quorum sensing system, another notable advantage of the Grant Library is that we selected for the transposon insertion and the plasmid that confers IPTG-inducible natural competence during outgrowth in selective media. Accordingly, upon induction with IPTG, mutants from the Grant Library can undergo additional genome edits when co-incubated with gDNA. Furthermore, growth in the absence of selection leads to rapid curing of the competence plasmid if it is not needed. These features make the Grant Library an invaluable resource for studying pleiotropic and epistatic genetic interactions in *V. cholerae*.

Indeed, the non-linearity between paired mutants when grown on inosine demonstrates this utility. Ongoing work in the Grant lab involves utilizing the two transposon libraries described in this work, and a third high cell density locked library we are generating, to deconstruct gene *x* transcriptional state *x* environmental interactions, including synthetically lethal genetic combinations. Such information will be invaluable toward generating new therapeutic options to treat *V. cholerae* infections.

Construction of the Grant Library was aided by a novel, in-house gDNA extraction method we developed based on thymol cell lysis. Our use of thymol allowed extraction of gDNA in a highly efficient and economical way. Indeed, N.A.G. has performed as many as 576 manual gDNA extractions in a single workday. Additionally, at the concentrations used in our studies, gDNA extracted with thymol does not negatively impact its use in other downstream applications, including PCR amplification, whole genome and Sanger sequencing. Given thymol’s cell killing efficacy in *Vibrio*, we tested the treatment on several other biological specimens including bacteria in gram -negative and -positive staining classes and yeast. Among our test strains, gDNA yield was highest for *V. cholerae* (∼ 22 ng/µl), although most strains yielded some gDNA (2 – 8 ng/µl) (data not shown). That *Vibrio* is highly susceptible to thymol treatment is an interesting avenue for further research as its understanding can provide additional insight into its antibacterial activity, potentially contributing to novel disease control strategies. In summary, the Grant Library complements the non-redundant library made in the quorum-incompetent strain, and the methods used to construct it, a valuable approach toward understanding *V. cholerae* evolutionary genetics.

## Acknowledgements

We thank Ewen Cameron and the John Mekalanos lab for constructing the 2008 *V. cholerae* non-redundant library, Vic DiRita for granting access to a copy of the collection, and Bonnie Bassler and Julie Valastyan for providing plates that were missing from the DiRita collection and the gain of function *luxOG319S* parent strain. We also thank Ankur Dalia for providing the plasmid conferring natural competence. This work was supported by National Institutes of Health Grants GM139537 and AI158433 to C.M.W and a NIGMS Diversity Supplement to N.A.G. Financial support was also received from The BEACON Center for the Study of Evolution in Action at Michigan State University (to N.A.G).

