## Supplementary material for "Deployment of a *Vibrio cholerae* ordered transposon mutant library in a quorum-competent genetic background": Supp F1-F3, Tables S1-S4

**Supplementary information text**

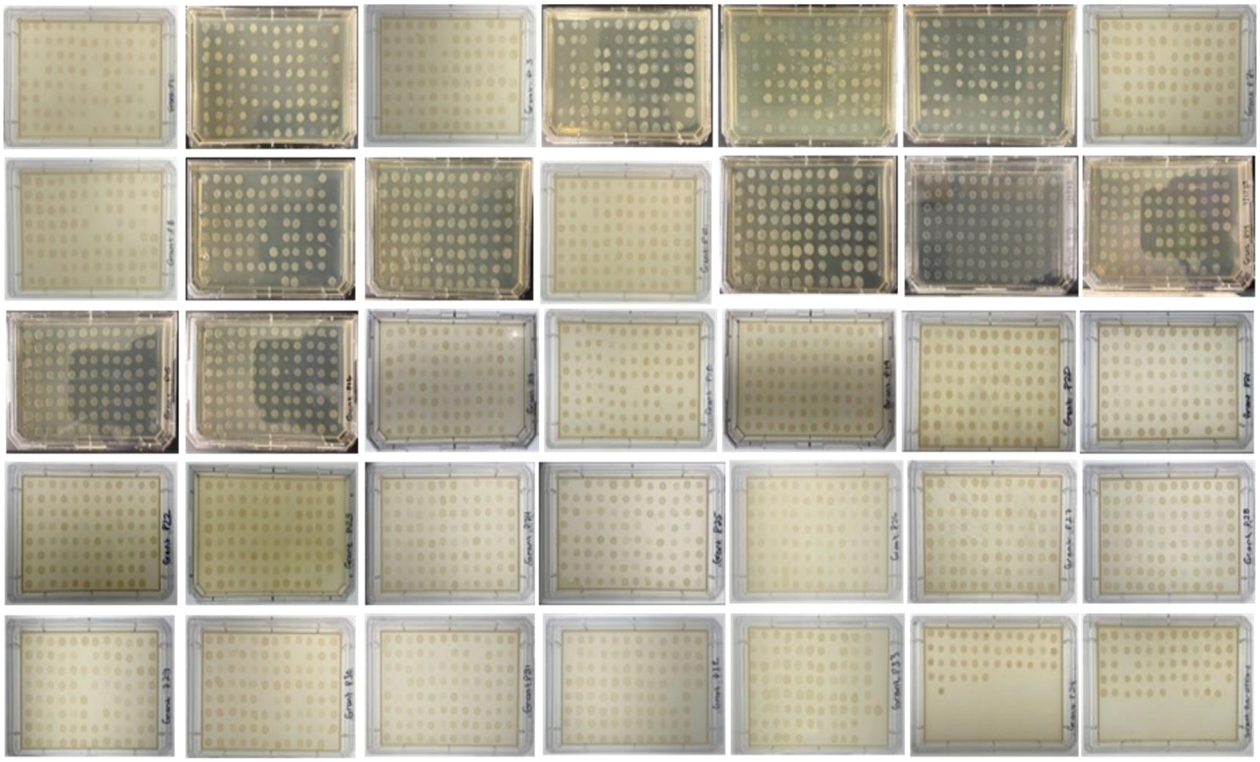

Figure S1. Growth of Grant library transposon mutants on antibiotic selective media. Each mutant was spot plated on LB agar + kanamycin and chloramphenicol to select for mutants with the transposon insertion and the plasmid conferring natural competence (pMMB-*tfoX*-*qstR*), respectively. Plates were incubated overnight, and the growth of each strain was recorded on the following day.

Figure S2 (panels A–S)

### Mutant 1D3

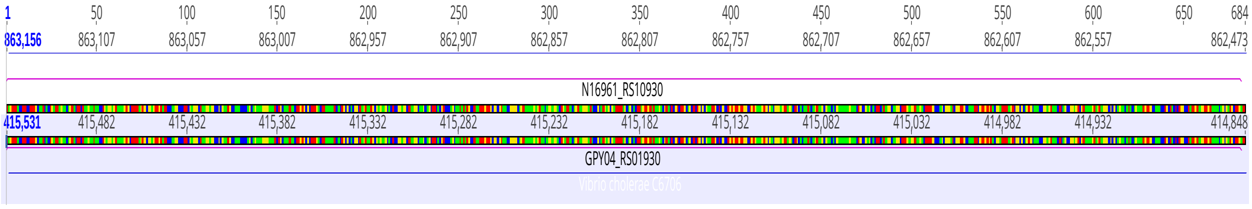


### Mutant 2C1

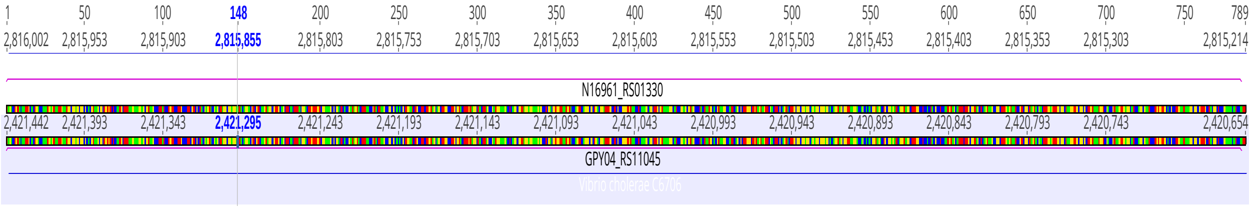


### Mutant 2D5

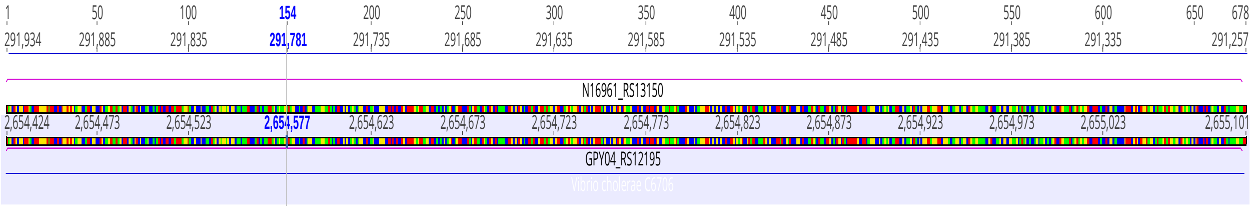


### Mutant 4B2

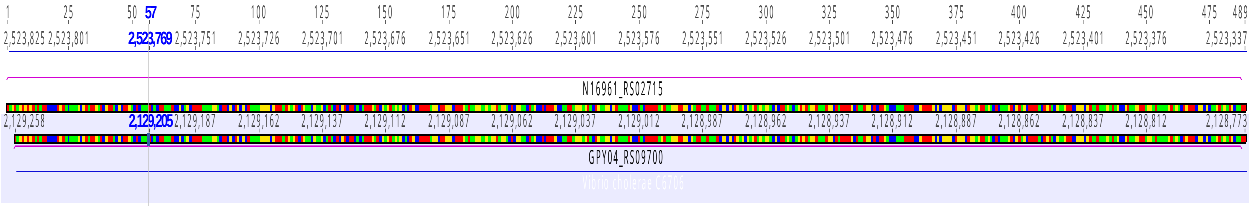

### Mutant 6H11

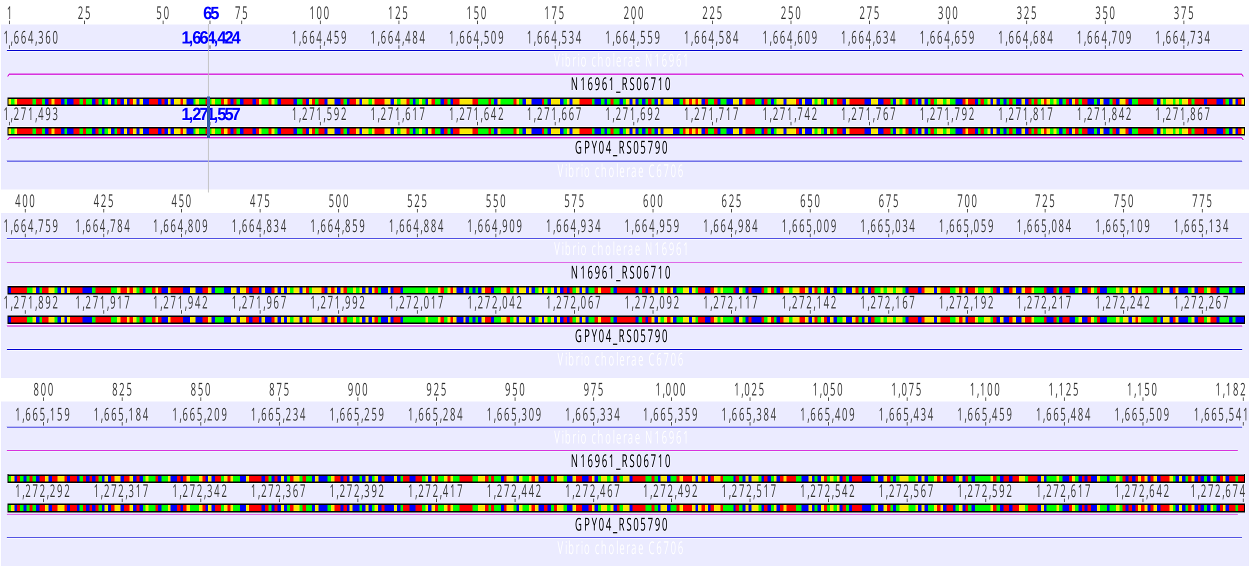


### Mutant 9E2

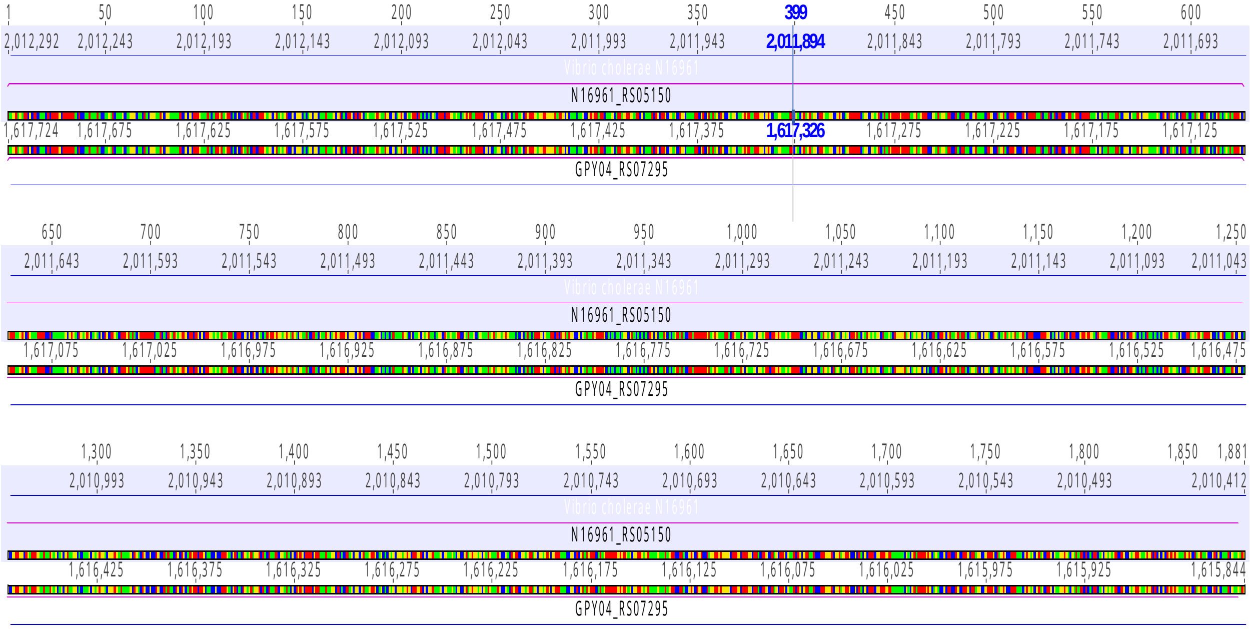

### Mutant 10H8

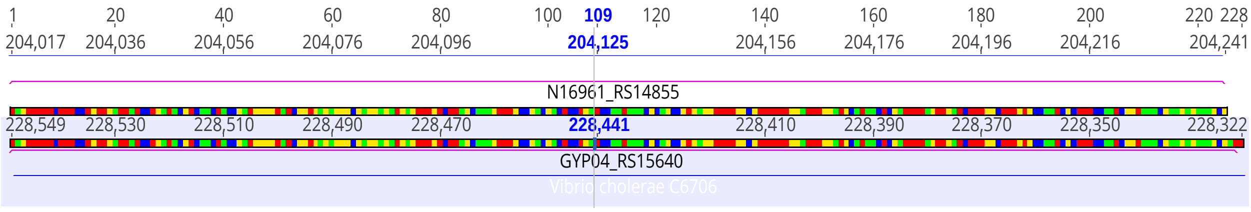


### Mutant 13H5

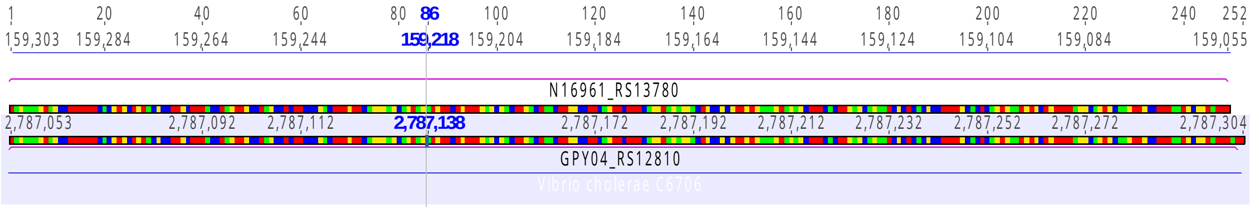


### Mutant 19H8

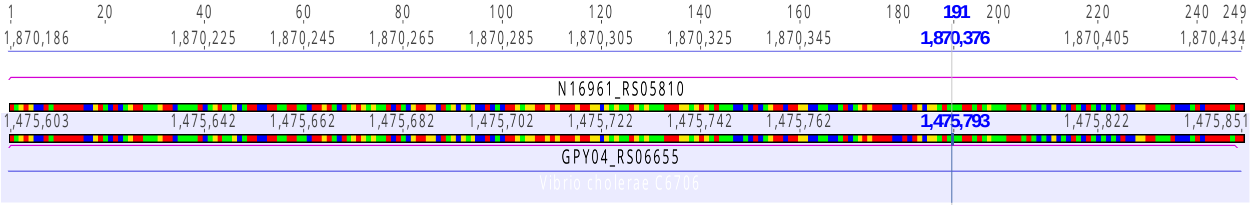


### Mutant 21C6

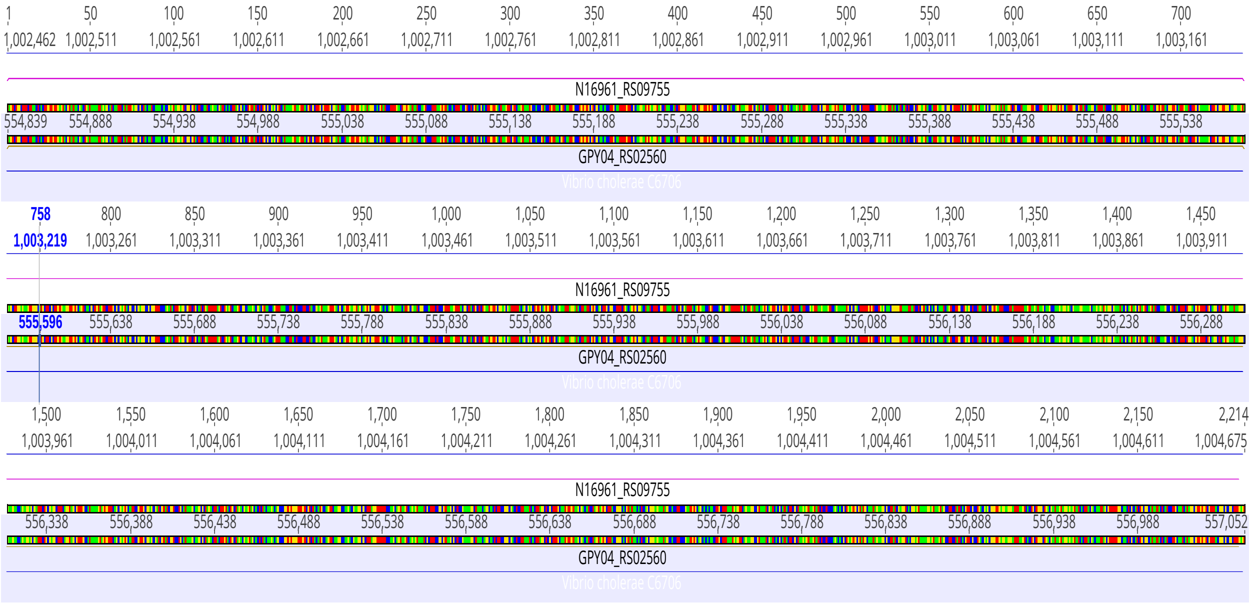

### Mutant 24B2

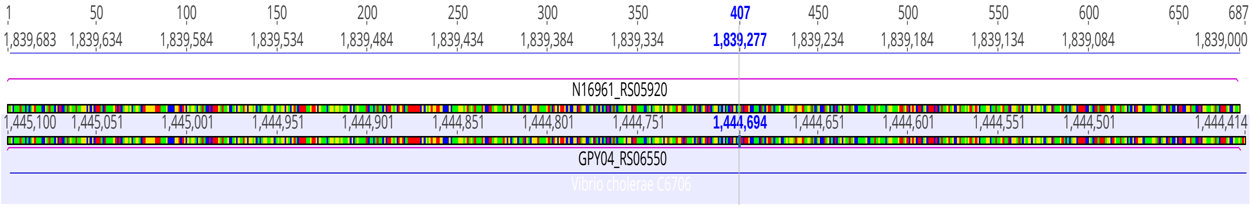


### Mutant 24D2

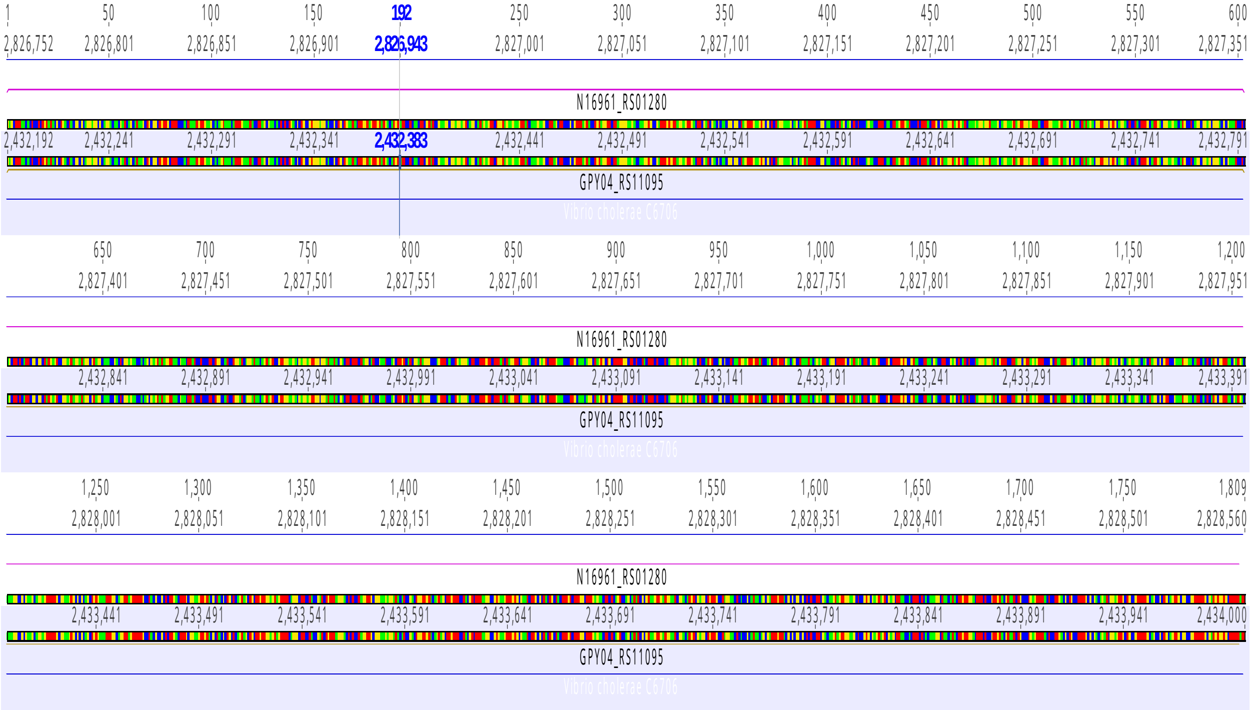


### Mutant 24G4

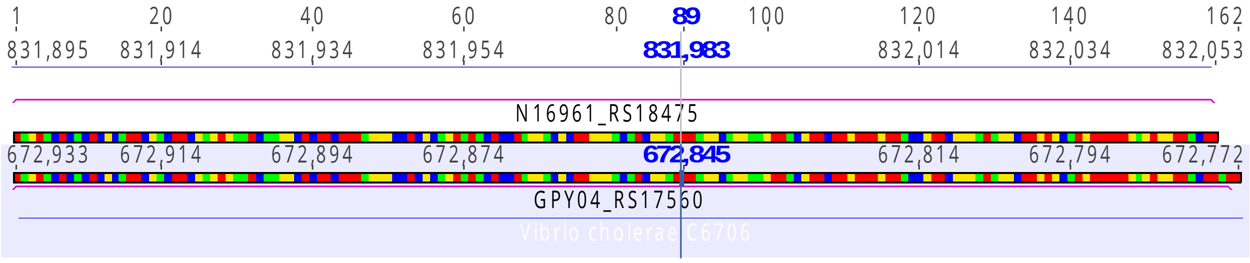

**Mutant 28E11**

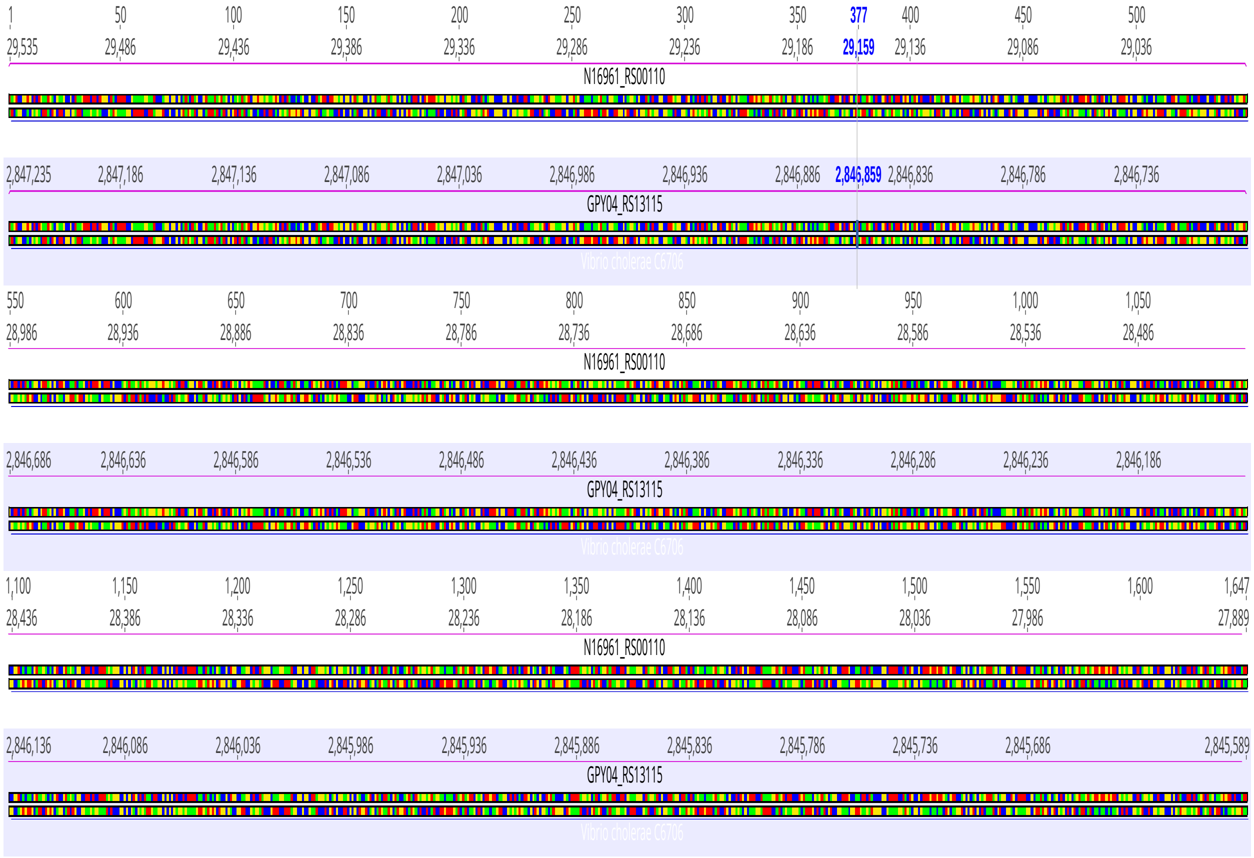

### Mutant 29E1

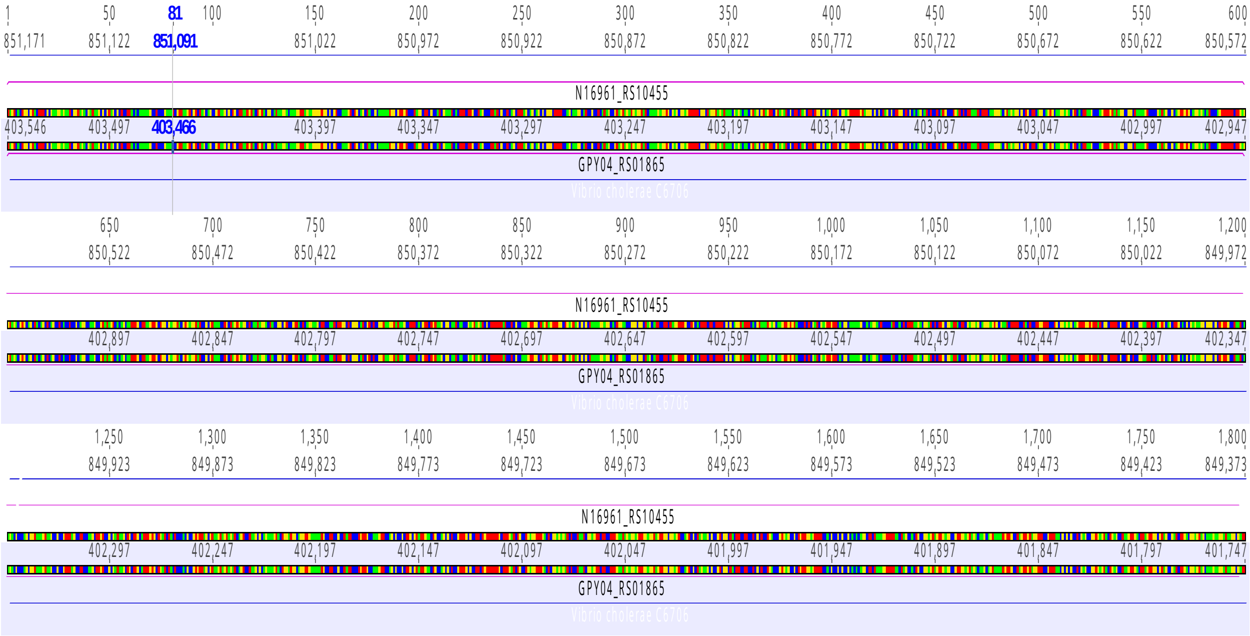


### Mutant 31A10

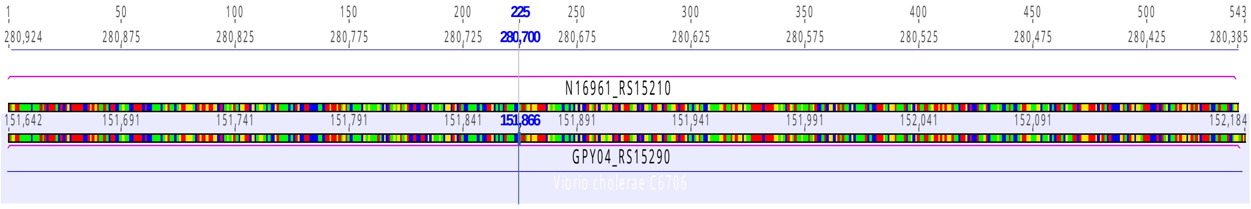


### Mutant 31E11

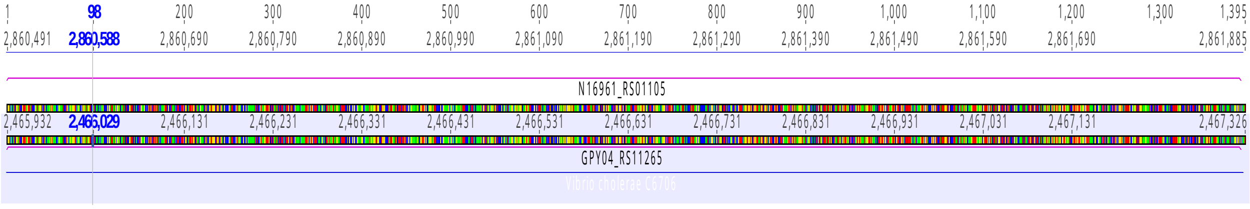

**Mutant 31F3**

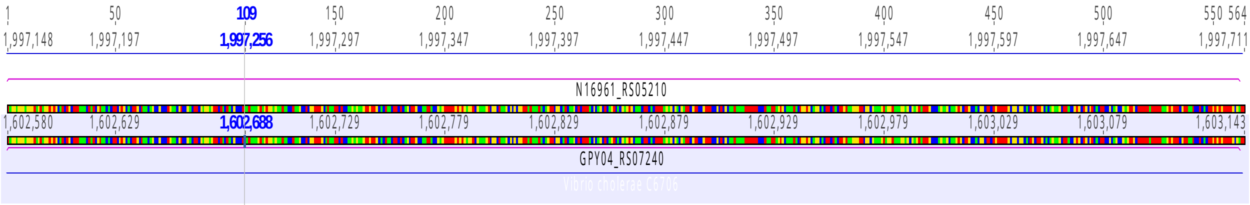


**Mutant 33C7**

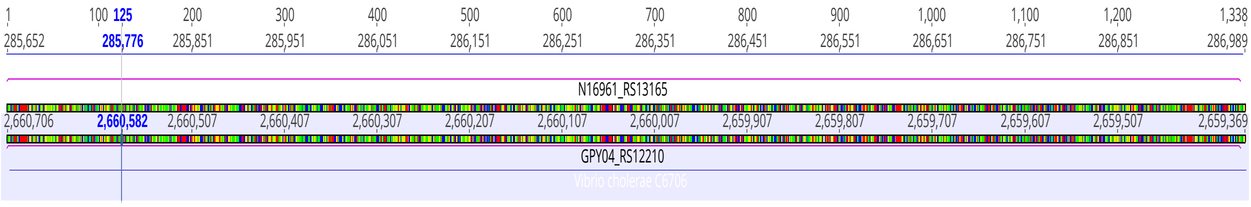


Figure S2: Comparison of Transposon Insertions between *V. cholerae* N16961 and C6706. Each panel (A–S) displays the alignment for genes that lie on the regression line (Fig. 5) in the indicated samples. Transposon insertion sites were determined from whole-genome sequence analysis of C6706 (this work) and for N16961 from Cameron et al. (2008). These sites are highlighted above the representation of each gene sequence in blue, and the names of each strain and the identity of each gene are shown within the colored blocks.

Figure S3 (panels A–D)

**Mutant 16B1**

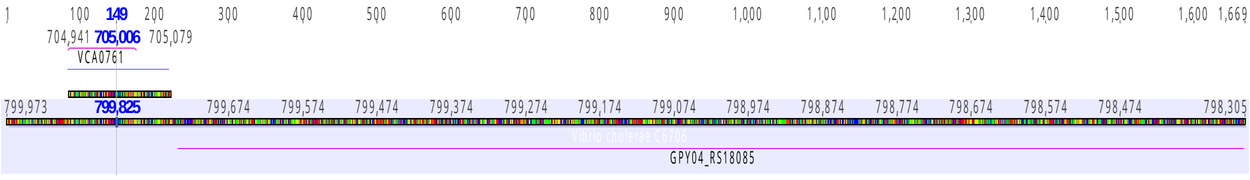


**Mutant 33F6**

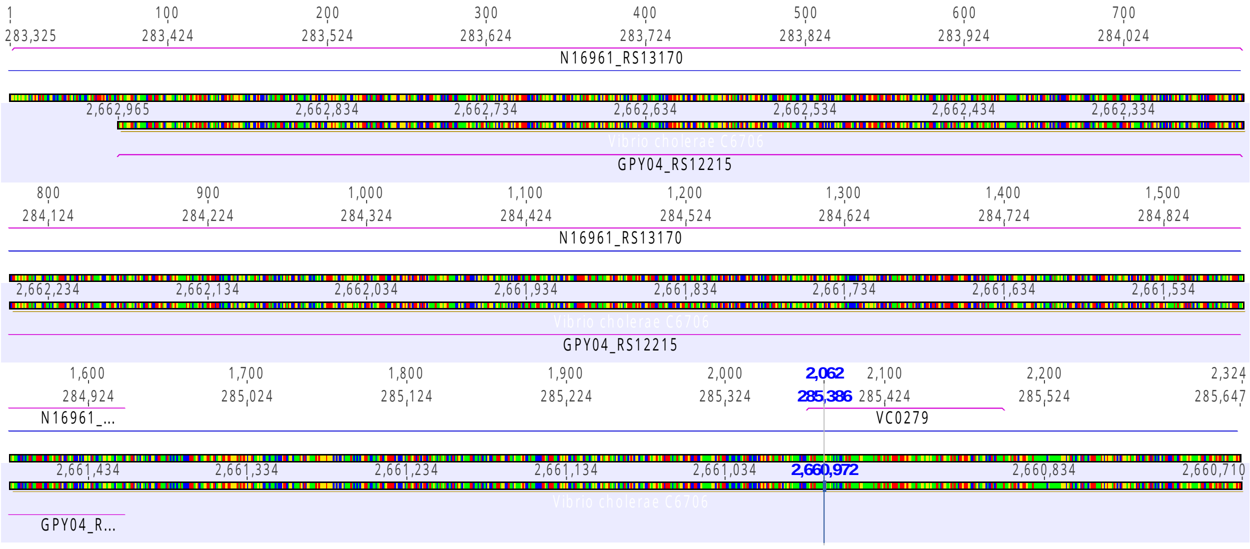

**Mutant 8E3**

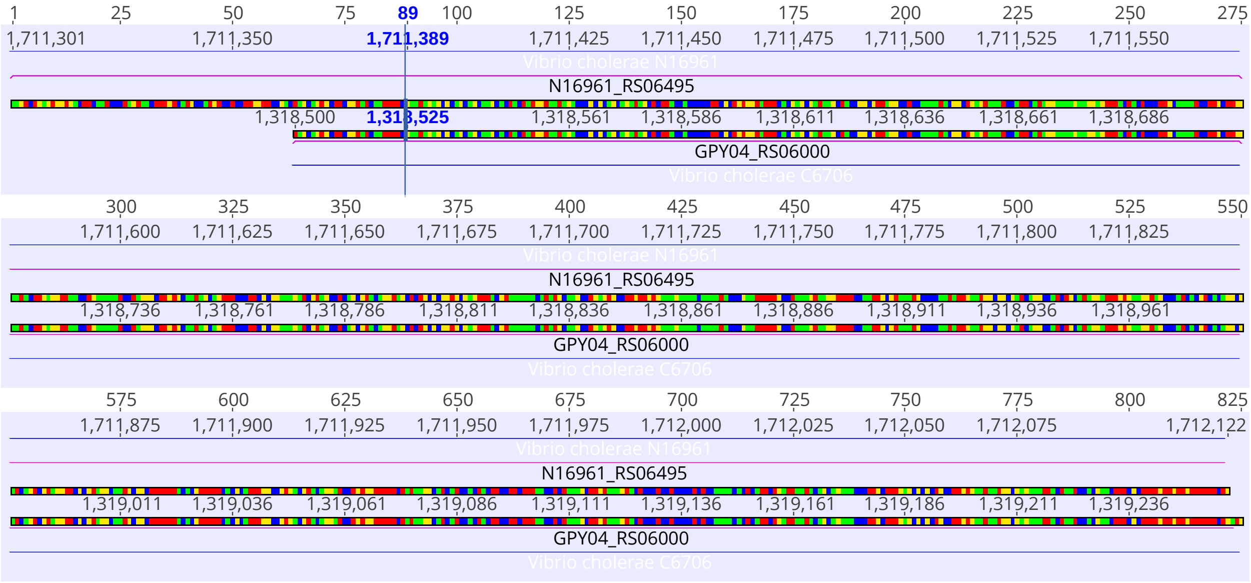


**Mutant 12D4**

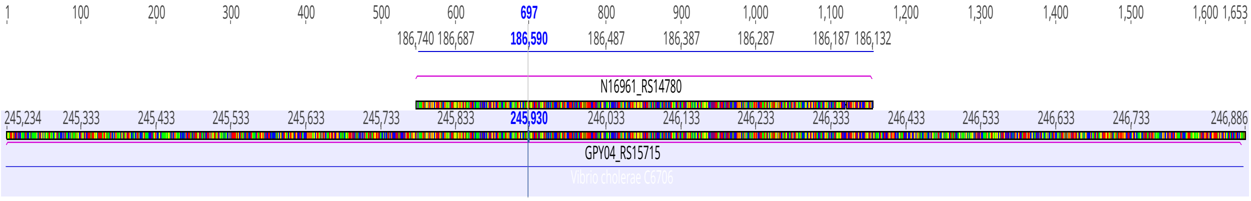


Figure S3: Comparison of Transposon Insertions between *V. cholerae* N16961 and C6706. Each panel (A–D) displays an alignment of genes whose points deviated from the regression line (Fig. 5) in the indicated samples. Panels A and B illustrate category one false negatives, where transposon insertions are mapped to genes annotated in *V. cholerae* N16961 but not in C6706. These correspond to points that fall on the y-axis in Fig. 5. Panels C and D illustrate category two false negatives, where each gene pair aligns with > 99.8% homology between N16961 and C6706, but they differ in size, accounting for the divergence seen for those points in Fig. 5.

Table S1. Gene identities for the bottom 2.5% of Grant library mutants grown in 20 mM inosine.

| Plate_index | Chromosome | Gene description | Gene name | Gene role |
|---|---|---|---|---|
| P26_A6 | 1 | Phosphoribosylformimino-5-aminoimidazole carboxamide ribotide isomerase | hisA | Amino acid biosynthesis |
| P15_F10 | 1 | Thiamin ABC transporter, periplasmic thiamin-binding protein | tbpA | Thiamin ABC transporter |
| P11_B10 | 1 | Imidazoleglycerol-phosphate dehydratase/histidinol-phosphatase | hisB | Amino acid biosynthesis |
| P26_C3 | 1 | Alanine dehydrogenase | Ald | Energy metabolism |
| P15_G12 | 1 | Cyclic AMP phosphodiesterase | cpdA | Regulatory functions |
| P19_G5 | 1 | thiH protein | thiH | Biosynthesis of cofactors, prosthetic groups, and carriers |
| P11_F6 | 1 | N-acetyl-gamma-glutamyl-phosphate reductase | argC | Amino acid biosynthesis |
| P11_A10 | 1 | Citrate synthase | gltA | Energy metabolism |
| P19_A2 | 1 | Indole-3-glycerol phosphate synthase/phosphoribosylanthranilate isomerase | trpC/F | Amino acid biosynthesis |
| P26_D10 | 1 | 3-dehydroquinate dehydratase | aroQ | Amino acid biosynthesis |
| P15_F3 | 1 | Cys regulon transcriptional activator | cysB | Regulatory functions |

Table S2. Gene identities for the bottom 2.5% of Cameron et al. library mutants grown in 20 mM inosine.

| Plate_index | Chromosome | Gene description | Gene name | Gene role |
|---|---|---|---|---|
| P26_A6 | 1 | Phosphoribosylformimino-5-aminoimidazole carboxamide ribotide isomerase | hisA | Amino acid biosynthesis |
| P15_G12 | 1 | Cyclic AMP phosphodiesterase | cpdA | Regulatory functions |
| P19_G5 | 1 | thiH protein | thiH | Biosynthesis of cofactors, prosthetic groups, and carriers |
| P11_A10 | 1 | Citrate synthase | gltA | Energy metabolism |
| P19_A9 | 1 | torD protein | torD | Unknown function |
| P11_F6 | 1 | N-acetyl-gamma-glutamyl-phosphate reductase | argC | Amino acid biosynthesis |
| P19_A2 | 1 | Indole-3-glycerol phosphate synthase/phosphoribosylanthranilate isomerase | trpC/F | Amino acid biosynthesis |
| P15_A10 | 1 | Sulfate ABC transporter, ATP-binding protein | cysA | Transport and binding proteins |
| P26_D10 | 1 | 3-dehydroquinate dehydratase | aroQ | Amino acid biosynthesis |
| P15_C5 | 1 | Serine acetyl transferase | cysE | Amino acid biosynthesis |
| P15_E5 | 1 | Flagellar rod protein, putative | FlaI | Cellular process |

Table S3. Gene identities for the top 2.5% of Grant library mutants grown in 20 mM inosine.

| Plate_index | Chromosome | Gene description | Gene name | Gene role |
|---|---|---|---|---|
| P11_D10 | 2 | Hypothetical protein | | Methyl-accepting chemotaxis protein |
| P26_E4 | 2 | Conserved hypothetical protein | | |
| P2_E3 | 1 | Flagellin FlaC | flaC | Cellular process |
| P19_D4 | 2 | Amino acid ABC transporter, permease protein; DUF3296 family protein (Geneious) | | Transport and binding proteins |
| P26_D6 | 1 | PTS system, N-acetylglucosamine-specific IIABC component | nagE | Transport and binding proteins |
| P26_D4 | 1 | Elongation factor G | fusA-2 | Protein synthesis |
| P26_F3 | 1 | Antioxidant, putative; Bifunctional molybdopterin-guanine dinucleotide biosynthesis adaptor protein MobB/molybdopterin molybdotransferase MoeA (Geneious) | MobB/MoeA | Cellular process |
| P26_B7 | 1 | Rfbl protein | rfbL | Unknown function |
| P19_D5 | 1 | CDP-ribitol pyrophosphorylase-related protein | thrA | Fatty acid and phospholipid metabolism; bifunctional aspartate kinase/homoserine dehydrogenase I |
| P19_C4 | 2 | Resolvase, putative; IS3-like element ISVch4 family transposase (Geneious) | | Mobile and extrachromosomal element functions |
| P11_G12 | 1 | Sensor histidine kinase; Outer membrane transport protein (Genious) | | Regulatory functions |
| P26_E8 | 1 | Syd protein | Syd | Unknown function |

Table S4. Gene identities for the top 2.5% of Cameron et al. library mutants grown in 20 mM inosine.

| Plate_index | Chromosome | Gene description | Gene name | Gene role |
|---|---|---|---|---|
| P26_F2 | 1 | Conserved hypothetical proteins; 2,3-dihydroxtbenzoate dehydrogenase (Geneious) | | Hypothetical protein |
| P26_C2 | 1 | Glutamate synthase, large subunit | gltB-1 | Amino acid biosynthesis |
| P26_C4 | 1 | ABC transporter, ATP-binding protein; transglycosylase domain-containing protein/penicillin binding protein (Geneious) | | Transport and binding proteins |
| P26_G7 | 1 | Hypothetical protein | | |
| P15_D5 | 1 | Conserved hypothetical protein; Bcr/CflA family multidrug efflux MFS transporter (Geneious) | | Hypothetical proteins |
| P2_E10 | 1 | Formate transporter 1, putative | trxB (Geneious) | Transport and binding proteins; thioredoxin-disulfide reductase activity |
| P11_E7 | 1 | Trypsin, putative; efflux RND transporter permease subunit (Geneious) | vexK (Geneious) | Protein fate |
| P26_D4 | 1 | Elongation factor G | fusA-2 | Protein synthesis |
| P11_D5 | 1 | A/G-specific adenine glycosylase | mutY | DNA metabolism |
| P11_E10 | 1 | Polysaccharide biosynthesis protein, putative; methylenetetrahydrofolate dehydrogenase (NADP+) activity (Geneious) | folD (Geneious) | Cell envelope |
| P19_H10 | 2 | Hypothetical protein; VOC family protein (Geneious) | | |

Dataset S1. Plate indices file

Dataset S2. Modified version of transposon library mutant dataset presented in Cameron et al., 2008.

Dataset S3. List of genes duplicated in the Cameron et al., 2008 transposon library mutant dataset.

Dataset S4. List of gene ortholog pairs present in *Vibrio cholerae* strains N16961 and C6706.

Dataset S5. List of genes unique to *Vibrio cholerae* strain N16961.

Dataset S6. List of genes unique to *Vibrio cholerae* strain C6706.

Dataset S7. List of transposon insertion mutants within a 50 kb window upstream and downstream of *luxO*.

Dataset S8. Sanger sequence chromatograms for 94 transposon mutants sequenced in 50 kb window upstream and downstream of *luxO*.

Dataset S9. $OD_{600}$ readings over time monitoring growth of Grant and Cameron et al. parent strains.

Dataset S10. Endpoint $OD_{600}$ readings for 5-paired plates from Grant and Cameron et al. transposon mutant libraries.

Table S1. Gene identities for the bottom 2.5% of Grant library mutants grown in 20 mM inosine.

Table S2. Gene identities for the bottom 2.5% of Cameron et al. library mutants grown in 20 mM inosine.

Table S3. Gene identities for the top 2.5% of Grant library mutants grown in 20 mM inosine.

Table S4. Gene identities for the top 2.5% of Cameron et al. library mutants grown in 20 mM inosine.
